# Integrating Multiplex SiMPull and Computational Modeling to Evaluate Combinatorial Aspects of EGFR Signaling

**DOI:** 10.1101/227512

**Authors:** Emanuel Salazar-Cavazos, Carolina Franco Nitta, Eshan D. Mitra, Bridget S. Wilson, Keith A Lidke, William S. Hlavacek, Diane S. Lidke

## Abstract

The Epidermal Growth Factor Receptor (EGFR/ErbB1/HER1) plays an important role in both physiological and cancer-related processes. To study the factors that influence receptor phosphorylation, we have coupled Single Molecule Pull-down (SiMPull) measurements with rule-based modeling of EGFR signaling. Using SiMPull, we quantified the phosphorylation state of thousands of individual receptors. These measurements enabled the first direct detection of multisite phosphorylation on full-length EGFR and revealed that the extent of phosphorylation varies by tyrosine site and is dependent on the relative abundance of signaling partners that limit access by tyrosine phosphatases. We also evaluated the impact of oncogenic mutations and ligands with varying affinity on phosphorylation kinetics. Simulations highlight the importance of dimer lifetimes on EGFR phosphorylation and signaling output.

## INTRODUCTION

The ability of a cell to respond rapidly and specifically to changes in the surrounding environment is controlled by protein-protein interactions at the plasma membrane and along the signaling cascade. While much is known about the biochemical events that govern signaling pathways, this information has mostly been derived from population-based measurements that average over millions of cells and/or proteins. However, there is growing evidence that system heterogeneity at both the cellular and molecular levels contribute to cellular information processing (Lahav et al. 2004; Feinerman et al. 2008; Coba et al. 2009; Spencer et al. 2009). EGFR has 20 cytoplasmic tyrosines, at least 6 of which (Y992, Y1045, Y1068, Y1086, Y1148, and Y1173) are capable of recruiting signaling proteins when phosphorylated (Jorissen et al. 2003; Schulze, Deng, and Mann 2005; Hause et al. 2012). The potential for multisite phosphorylation on individual receptors provides a mechanism to control signaling output that is dependent on complex factors such as steric hindrance, relative abundance of receptors and adaptors, cooperative interactions and lifetimes of complexes (Gibson, Parkes, and Liebman 2000; Salazar and Höfer 2009; Coba et al. 2009; Lau et al. 2011; Stites et al. 2015). Signal propagation is opposed by cellular phosphatases (Kleiman et al. 2011), whose activity is proposed to be blocked when phosphotyrosine substrates are engaged with binding partners (Rotin et al. 1992; Brunati et al. 1998; Jadwin et al. 2018). To better understand the contributions of factors such as adaptor abundance and phosphatase access on protein phosphorylation heterogeneity, we have combined quantitative single molecule measurements with rule-based modeling of EGFR signaling.

SiMPull is a powerful technique that allows for interrogation of macromolecular complexes at the individual protein level. This technique captures macromolecular complexes on glass coverslips, combining the power of immunoprecipitation with high resolution imaging for single molecule quantitative analysis (Jain et al. 2011). Here, we describe improvements to the SiMPull method that enable accurate evaluation of phosphorylation at multiple sites on individual receptors. This represents a significant improvement over traditional, semi-quantitative methods such as Western Blot analysis or flow cytometry, which report only trends in phosphorylation state changes over the entire population. Our improvements to SiMPull protocols include pretreatment to reduce autofluorescence and corrections for receptor surface expression. We employed a simplified imaging chamber that accommodates up to 20 samples, each with sample volumes of only 10 μl. We also demonstrate the critical importance of optimizing antibody labeling and fixation conditions. To quantify receptor phosphorylation, we have used two- and three-color imaging to identify individual proteins and their corresponding phosphorylation status. These multiplex SiMPull measurements provide a new level of detail in the status of receptor phosphorylation that was previously inaccessible with traditional biochemical techniques.

The unique data provided by SiMPull was used to parameterize a mathematical model for site-specific phosphorylation kinetics of EGFR and to test model predictions. Particularly, we explored two traits of relevance for signaling processes: 1) the influence of adaptor protein abundances on the phosphorylation levels of the EGFR residues to which these proteins bind; and 2) the frequency of multi-site phosphorylation across individual EGFR in the population. We specifically explore how the phosphorylation status of individual receptors reflects the opposing contributions of adaptor binding versus phosphatase activity. For example, simulations predicted that overexpression of the adaptor protein Grb2 would lead to increase in phosphorylation at the EGFR tyrosine residue where Grb2 binds (Y1068). This prediction was confirmed experimentally using SiMPull and compared to Shc1 binding Y1173. Additionally, our model predicts that differences in relative adaptor protein abundances across different cell lines should result in different phosphorylation patterns, consistent with phosphoproteome and total protein expression profiles reported by Shi et al. (Shi et al. 2016). Our model also uniquely evaluates the experimentally observed phosphorylation kinetics of mutant versus wildtype EGFR, as well as EGFR engaged by the low-affinity ligand epigen (EPGN) versus the high-affinity ligand EGF. Simulation results suggest that the differential phosphorylation patterns are explained best by unique dimer lifetimes. These new insights into the extent of phosphorylation at individual tyrosines along with the existence of multisite phosphorylation has implications for how EGFR translates extracellular cues into downstream signaling outcomes.

## RESULTS

### Assessing receptor phosphorylation at the single molecule level

Our principal method to interrogate the phosphorylation status of individual EGFR after different treatments is single molecule immunoprecipitation via SiMPull. SiMPull samples are prepared in a manner similar to SDS-PAGE/Western Blot protocols, but the sample is evaluated using single molecule microscopy to dramatically improve quantification. Figure 1A illustrates the basic principles of our assay. Briefly, cells expressing GFP-tagged EGFR are lysed before or after treatments (as specified in figure legends). Clarified lysates are dispensed onto coverslips sparsely pre-coated with anti-EGFR antibodies. Following incubation and washes, individual EGFR-GFP are imaged by Total Internal Reflection Fluorescence (TIRF) microscopy. In the raw images, receptors appear as diffraction limited fluorescent spots that may represent more than one molecule (Fig. 1B, top left). Single molecules are selected by fitting each emission profile to a point spread function model and rejecting fluorescent spots that do not fit to a single emitter model (Fig. 1B, bottom left). To evaluate the overall phosphorylation state of individual receptors, coverslips are incubated with pan-reactive, anti-phosphotyrosine (anti-PY) antibodies bearing fluorescent tags with spectral emission distinct from GFP (Fig. 1B, center). Colocalization between EGFR-GFP and anti-PY identifies phosphorylated receptors. In this example, the overlay image shows that 34.4% of EGFR-GFP are phosphorylated on at least one tyrosine site (Fig. 1B, right) Phosphorylation at specific tyrosines in the EGFR cytoplasmic tail is evaluated by substituting a site-specific phosphotyrosine antibody (anti-pY1068 or anti-pY1173) for the pan-PY antibodies.

**Figure 1.**
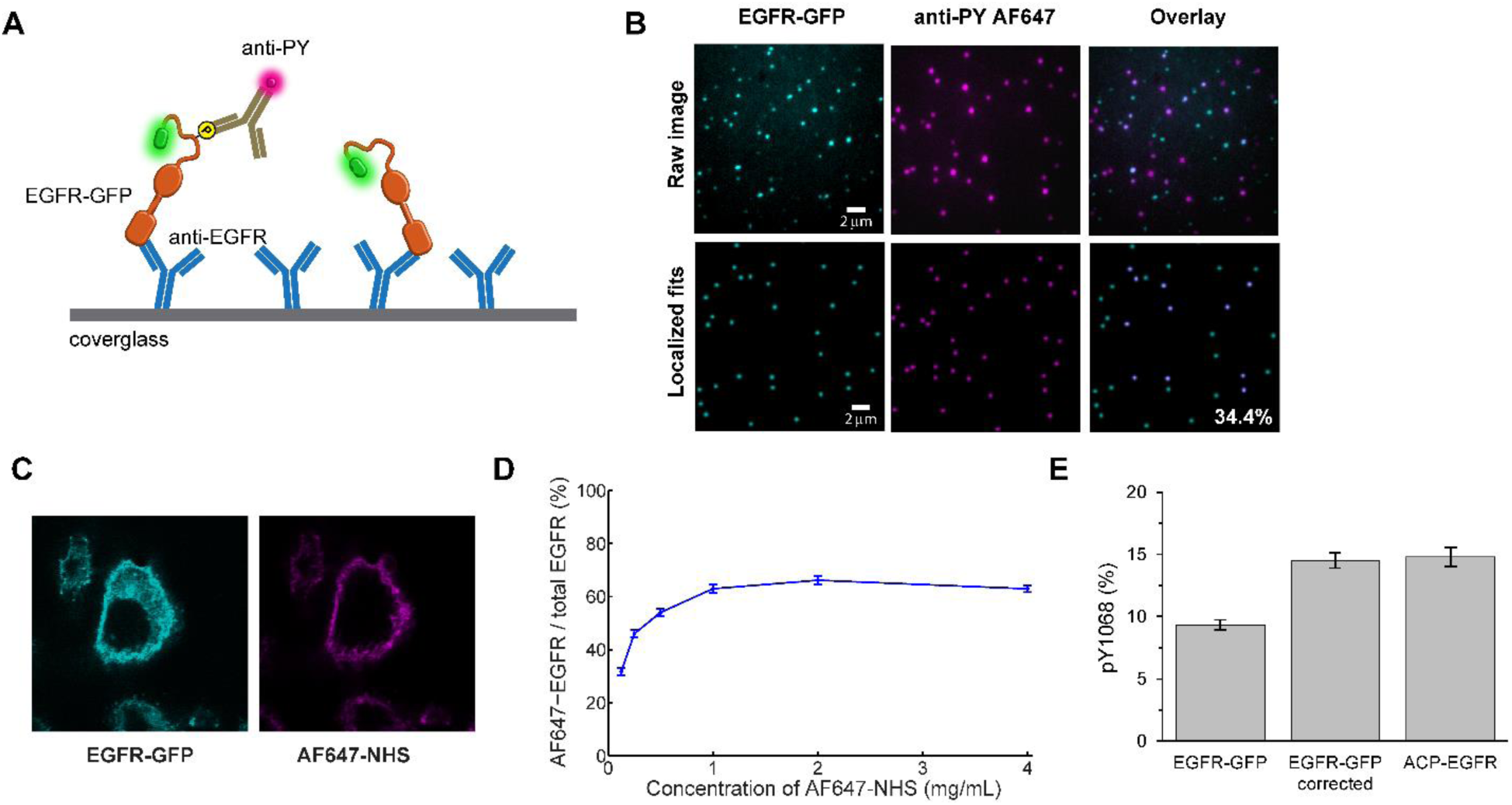
SiMPull to quantify protein phosphorylation. **(A)** Illustration depicting overall principle for assessing phosphorylation at the single molecule level using GFP-tagged EGFR (EGFR-GFP) as an example. **(B)** Representative images showing raw data (top) and blob-reconstructed localized molecules (bottom). CHO-EGFR-GFP cells were stimulated for 5 min with 25 nM EGF at 37°C before lysis for SiMPull. Raw images are brightness and contrast enhanced for visualization. The EGFR-GFP fits were filtered based on their fit to the microscope point spread function and the GFP-channel used as a mask to create the overlay. The number in the bottom right image represents the phosphorylation percentage estimated for this field of view. (**C)** Confocal images showing typical distribution of EGFR-GFP in CHO cells (left) and the labeling of surface proteins achieved with the AF647-NHS ester (right). **(D)** Cells were incubated with increasing concentrations of AF647-NHS and assayed by SiMPull to determine the percentage of EGFR-GFP molecules labeled with AF647. Number of receptors analyzed per data point, 850 < N < 1550. **(E)** Percentage of pY1068+ receptors estimated for EGFR-GFP before and after correcting for surface expression. The corrected phosphorylation percentage for EGFR-GFP corresponds to the value measured for ACP-EGFR, which only includes plasma membrane localized receptors. N>2400 for each EGFR type. Error bars are standard error of measured phosphorylation percentages.

Over the course of this study, we implemented a series of improvements in the experimental process and image analysis to achieve efficient and accurate quantification. These are described briefly here and more detail is found in Materials and Methods and in legends to Supplemental Figures S1-3. As has been previously noted (Jain et al. 2012), autofluorescence background was observed in our green spectral channel (503-548 nm) that was identified as single GFP molecules in the absence of cell lysate. We found that incubating the PEG-coated coverglass with Sodium Borohydride (10 mg/mL NaBH_4_ for 4 min) significantly reduced the number of background fluorescent molecules (Supplemental Fig. S1A, B). To increase our throughput, we generated a simple sample chamber by using a hydrophobic barrier pen to create an array of up to 20 independent regions on a coverglass that require as little as 10 µL of sample (Supplemental Fig. S1C). Since antibodies are used to quantify protein phosphorylation, it is critical to optimize the antibody labeling conditions, including time and concentration (Supplemental Fig. S1D-F). We found that on the time scale of our image acquisition the antibody could dissociate from EGFR (Supplemental Fig. S1E) which would lead to an underestimate of receptor phosphorylation. Post-antibody labeling fixation with 4% Paraformaldehyde/0.1% Glutaraldehyde (PFA/GA) for 10 min stabilized the antibody levels for at least 1 hr (Supplemental Fig. S1E, PFA/GA). Since binding affinity will vary for each antibody and fluorescent-conjugation may also alter antibody affinity, it is necessary to perform a binding curve for each antibody (Supplemental Fig. S1F) to ensure optimal labeling conditions.

### Correcting for EGFR Surface Expression in SiMPull Data Analysis

Like all immunoprecipitation methods that rely on detergent permeabilization of cells as a first step, SiMPull does not distinguish between EGFR trafficking through different intracellular compartments. We observed intracellularly-localized EGFR-GFP (Fig. 1C, left) that would be inaccessible to ligand. To determine the fraction of EGFR at the plasma membrane, we labeled all surface proteins on the CHO-EGFR-GFP cells with membrane-impermeable AF647-NHS Ester (Fig. 1C, right) and used SiMPull to visualize the amount of EGFR-GFP colocalized with AF647. By increasing the concentration of AF647-NHS until saturation is achieved, we estimated that ∼65% of the receptors are located at the plasma membrane (Fig. 1D). With this information, we corrected our measurements to account for only those receptors available to bind ligand (see Methods). For example, application of this correction shows that ∼14% of the receptors are phosphorylated at Y1068 after 1 min stimulation with 50 nM EGF (Fig. 1E). We note that while surface labeling of receptors with AF647-NHS ester allows for identification of surface proteins, we found that pre-labeling of EGFR in this way reduced EGF binding (data not shown). Therefore, we did not use global NHS-labeling of receptors for the study of EGFR activation. To validate our correction method, we analyzed the phosphorylation levels of receptors from CHO cells expressing ACP-tagged EGFR. We directly labeled the plasma membrane localized EGFR using membrane-impermeable CoA-Atto488 as described previously (Valley et al. 2015; Ziomkiewicz et al. 2013). Cells were then exposed to EGF and probed for EGFR phosphorylation with SiMPull, this time using Atto488 as the marker for plasma membrane EGFR. The percent of phosphorylated EGFR was similar when comparing the membrane-localized ACP-EGFR and the membrane-corrected EGFR-GFP samples (Fig. 1E). Therefore, the effects of EGF binding to EGFR on the plasma membrane can be accurately determined from whole cell lysates and we apply this correction for the remainder of the results.

### Extent of phosphorylation varies by tyrosine residue

Results of the use of SiMPull to characterize phosphorylation of EGFR in CHO-EGFR-GFP cells over a range of EGF doses are shown in Figure 2A. The multi-well hydrophobic array format made it possible to efficiently examine a full dose response of activation in a single imaging session. We quantified total EGFR tyrosine phosphorylation (PY) and compared it with the phosphorylation patterns for two specific tyrosine sites (Y1068 and Y1173). Cells simulated for 5 min (Fig. 2A) with increasing concentrations of EGF showed the expected increase in total phosphorylation with ligand dose (Fig 2A, PY, blue bars). This fraction reached 40.8 +/−1.3% with 50 nM EGF, a dose that is considered saturating. While the extent of phosphorylation on specific tyrosines are lower than total PY values across the dose response curve, the fraction of EGFR with phosphorylation at Y1173 was consistently higher than at Y1068 (Fig. 2A).

**Figure 2.**
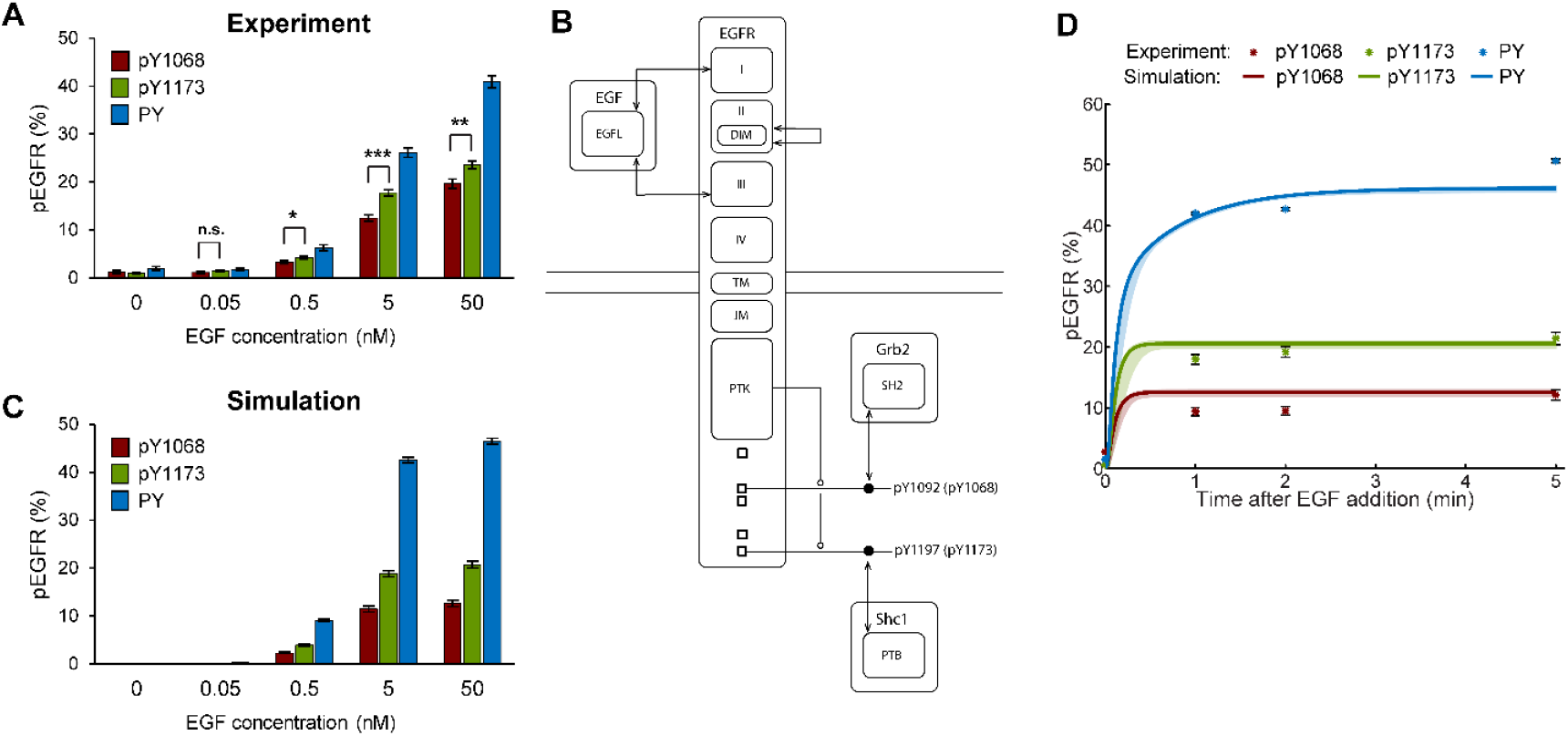
The extent of phosphorylation varies by tyrosine residue. **(A)** Dose response curve determined by SiMPull measurements for CHO-EGFR-GFP cells after 5 min of EGF addition at 37°C. Number of receptors analyzed per condition, N>1500. * P ≤ 0.05, ** P ≤ 0.01, *** P ≤ 0.001. **(B)** Schematic of EGFR tyrosine site-specific model, represented as an extended contact map (Chylek et al. 2011). **(C)** Simulation of dose response curve recapitulates the differences in extent of site-specific phosphorylation observed experimentally in (A). **(D)** Phosphorylation time course for CHO-EGFR-GFP cells stimulated with 25 nM EGF at 37°C. N>1800. Symbols are the SiMPull data and error bars are standard error of measured phosphorylation percentages. Solid lines are the model fit to the data points and the shaded areas are the prediction uncertainties with 95% confidence obtained through Bayesian uncertainty quantification (see Methods).

These results provide important observations. First, phosphorylation detection by SiMPull is sensitive, capable of detecting receptor phosphorylation at low ligand dose. Second, even under saturating ligand conditions, only a fraction of receptors is found to be phosphorylated, reaching a maximum of approximately 41% with 5 min stimulation. Third, the extent of phosphorylation varies by tyrosine residue. The detected phosphorylation levels are not restricted due to limitations in antibody labeling, since cells stimulated in the presence of phosphatase inhibitors showed increased receptor phosphorylation (Supplementary Fig. S2A,B) for all three antibodies. The addition of high salt (500 mM NaCl) in the buffer for the immunoprecipitation protocol, which interferes with electrostatic protein-protein interactions, did not change the detected phosphorylation (Supplementary Fig. S2C). This control suggests that, under our conditions, adaptor proteins that might interfere with antibody recognition in the SiMPull protocol are not co-precipitating.

### Computational model suggests variations in adaptor proteins abundances explain biased phosphorylation

To interpret the SiMPull data, we built a mathematical model based on our understanding of EGFR signaling mechanisms. We defined the model in terms of formal rules for interactions using the BioNetGen language (Faeder, Blinov, and Hlavacek 2009). Supplementary Table S1 summarizes key references and measurements that provide estimates of model parameters, including our own estimates of receptor and adaptor protein expression levels in the cell lines utilized in the study. Our model explicitly incorporates the asymmetric orientation of the EGFR kinase domains during dimerization, which assumes one monomer is active while the other monomer serves as an allosteric activator over the lifetime of that dimer pair (Zhang et al. 2006; Pryor et al. 2013). We also incorporated the observation that structural accessibility limits the efficiency of self-phosphorylation (i.e., within an active monomer’s own tail) by about 30% (Kovacs et al. 2015). An electronic version of the model is provided in the form of a BNGL file in the Online Supplementary Material (Supplementary File S1). Included with the BNGL file are all of the files necessary to perform the fitting and Bayesian uncertainty quantification procedures (Supplementary File S2). These files are also available online at the RuleHub repository [https://github.com/RuleWorld/RuleHub/tree/2019Jun18/Published/Salazar-Cavazos2019].

Figure 2B illustrates the rule-based structure of our model. Our first goal for computational modeling was to explore the possible mechanisms giving rise to the observed biased phosphorylation in EGFR at Y1173 compared to Y1068. As depicted in Figure 2B, and described in detail in Methods, our model includes site-specific phosphorylation of Y1068 and Y1173, and the recruitment of adaptor proteins Grb2 and Shc1 to these sites, respectively. We considered a range of parameter values in the model that might be adjusted to reproduce the biased phosphorylation. First, we considered the possibility that the phosphorylation and/or dephosphorylation rates are not equivalent for each tyrosine. While it is possible to fit our SiMPull data with this modification, differences in these rates are not supported by experimental measurements (see references in Supplementary Table S2). Thus, differences in these rates do not present a satisfactory explanation to account for the biased phosphorylation observed in our experiments.

An alternative possibility is that the binding of adaptor proteins (i.e., Grb2 and Shc1) physically serves as a barrier to protect specific phosphorylated tyrosines from phosphatase activity. This hypothesis is supported by *in vitro* and cellular studies showing the ability of SH2 domains to protect phospho-sites from dephosphorylation (Rotin et al. 1992; Brunati et al. 1998; Jadwin et al. 2018). To test if variation in protein abundances alone could explain our observations, we allowed the concentration of Grb2 and Shc1 to independently vary during the fitting process, while all other parameters were held constant. As a simplification, we took binding parameters (forward and reverse rate constants) for Grb2 interaction with pY1068 (via Grb2’s SH2 domain) and Shc1 interaction with pY1173 (via Shc1’s PTB domain) to be identical. The equilibrium dissociation constants measured *in vitro* by fluorescence polarization for these two interactions are comparable (2.6 uM for Grb2 SH2-pY1068 and 1.4 uM for Shc1 PTB-pY1173) (Hause et al. 2012). By introducing differential abundance of adaptor proteins, the model was able to recapitulate the time courses and dose responses determined by SiMPull, supporting the feasibility of this mechanism in generating biased phosphorylation (Figure 2C,D).

### Predicted influence of Grb2 overexpression in phosphorylation levels is observed experimentally

Our next goal was to experimentally validate the model prediction that biased phosphorylation is best explained by having different expression levels of Grb2 and Shc1. To set up conditions for these experiments, we ran simulations over a range of Grb2 overexpression values. Results in Figure 3A predict that Grb2 overexpression will lead to increased phosphorylation at Y1068, where Grb2 binds. Notably, no change is predicted for phosphorylated Y1173, where Grb2 is not expected to bind (Fig. 3B). To test these predictions, we created CHO-EGFR-GFP cells overexpressing Grb2-mCherry and evaluated the influence on EGFR phosphorylation using SiMPull (Fig. 3C,D). Consistent with simulation results, overexpression of Grb2 led to a marked increase in the phosphorylation of Y1068 (Fig. 3C). The relatively small change in phosphorylation at Y1173 (Figure 3D) is consistent with other reports that high expression of Grb2 can enhance phosphorylation at non-canonical binding sites (Jadwin et al. 2018).

**Figure 3.**
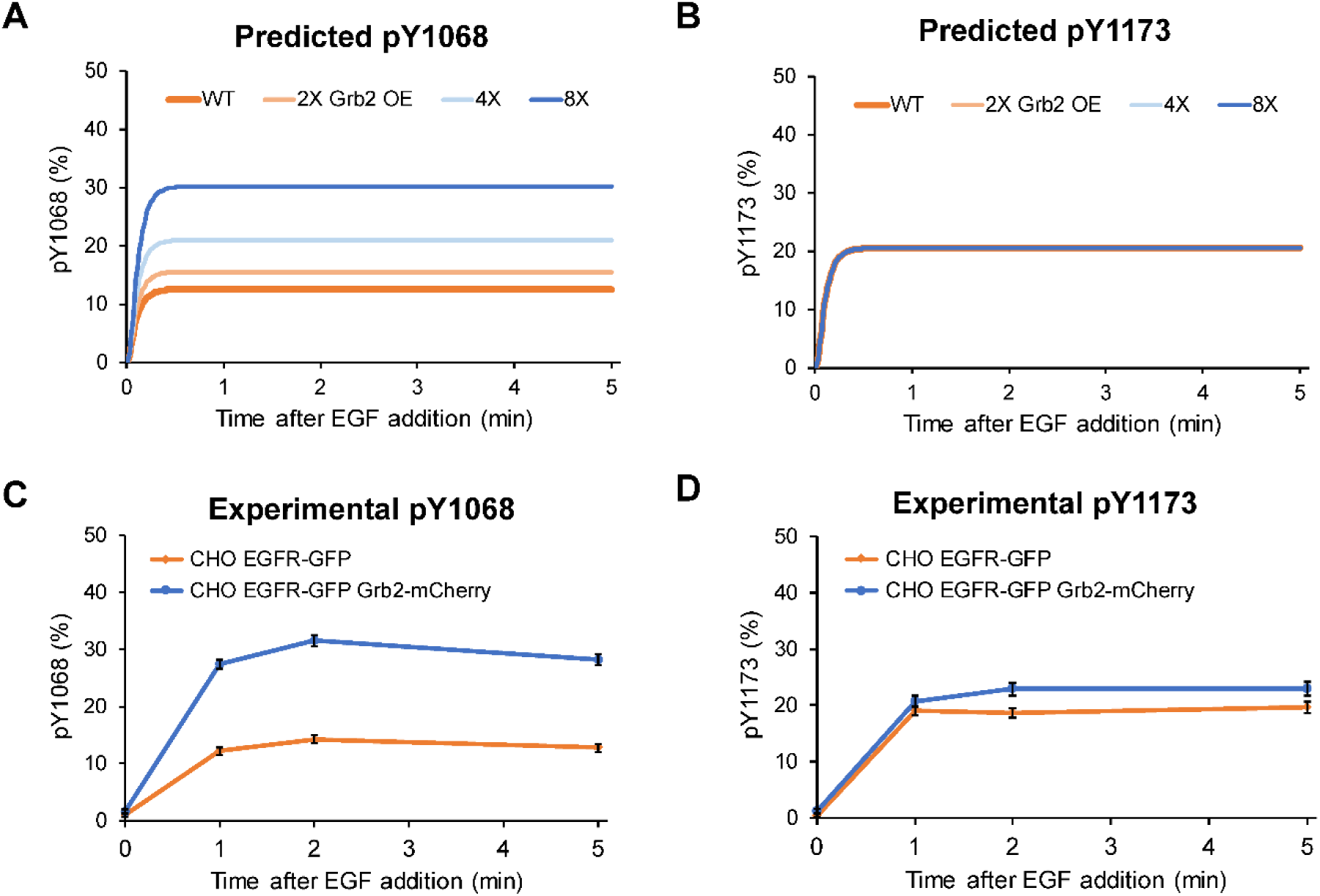
Predicted and observed phosphorylation kinetics in cells overexpressing Grb2. Predicted percentages of EGFR phosphorylation at tyrosines 1068 **(A)** and 1173 **(B)** after stimulation with 25 nM of EGF in cells with increasing overexpression (OE) of Grb2. SiMPull quantification of EGFR phosphorylation at tyrosines 1068 **(C)** and 1173 **(D)** after stimulation with 25 nM of EGF in CHO-EGFR-GFP cells expressing endogenous levels of Grb2 (orange) or overexpressing Grb2-mCherry (blue). N>1050. This cell line overexpresses Grb2 by approximately 3 fold on average, as determined by Western Blot analysis (data not shown). Error bars represent mean +/−S.E.M.

### Model predicts cell-specific phosphorylation patterns based on differences in adaptor protein abundances

Based on these results, we hypothesized that cell types naturally expressing different levels of these adaptor proteins would display different phosphorylation patterns. Protein copy numbers have been assayed using both global and targeted mass spectrometry-based proteomics in various cell lines (Shi et al. 2016; Kulak et al. 2014). These estimates include the protein copy numbers (per cell) for EGFR, Grb2 and Shc1 in the non-tumorigenic mammary epithelial HMEC and MCF10A cells, as well as in HeLa cervical cancer cells (see Supplementary Table S1). We performed simulations using these values and obtained model predictions for the phosphorylation patterns and kinetics in these three cell lines (Fig. 4A-C). In the HMEC cells, where the estimated abundances of both adaptor proteins are relatively low the model predicts similar levels of phosphorylation at both tyrosine residues (Fig. 4A). For the MCF10A cells the model predicts slightly higher phosphorylation at Y1173 given that its binding partner Shc1 is expressed in higher amounts than Grb2 (Fig. 4B). The most evident difference in expression levels between these adaptor proteins is found in HeLa cells, where it is estimated that there are ∼600,000 molecules of Grb2 per cell, compared to ∼100,000 molecules of Shc1 per cell. The model predicts that phosphorylation at Y1068 would be significantly higher than at Y1173 (Fig. 4C). Experimental results using SiMPull are shown in Figure 4D, confirming that HeLa S3 exhibit biased phosphorylation that is the reverse of that observed in CHO cells, with phosphorylation of Y1068 higher than phosphorylation at Y1173.

**Figure 4.**
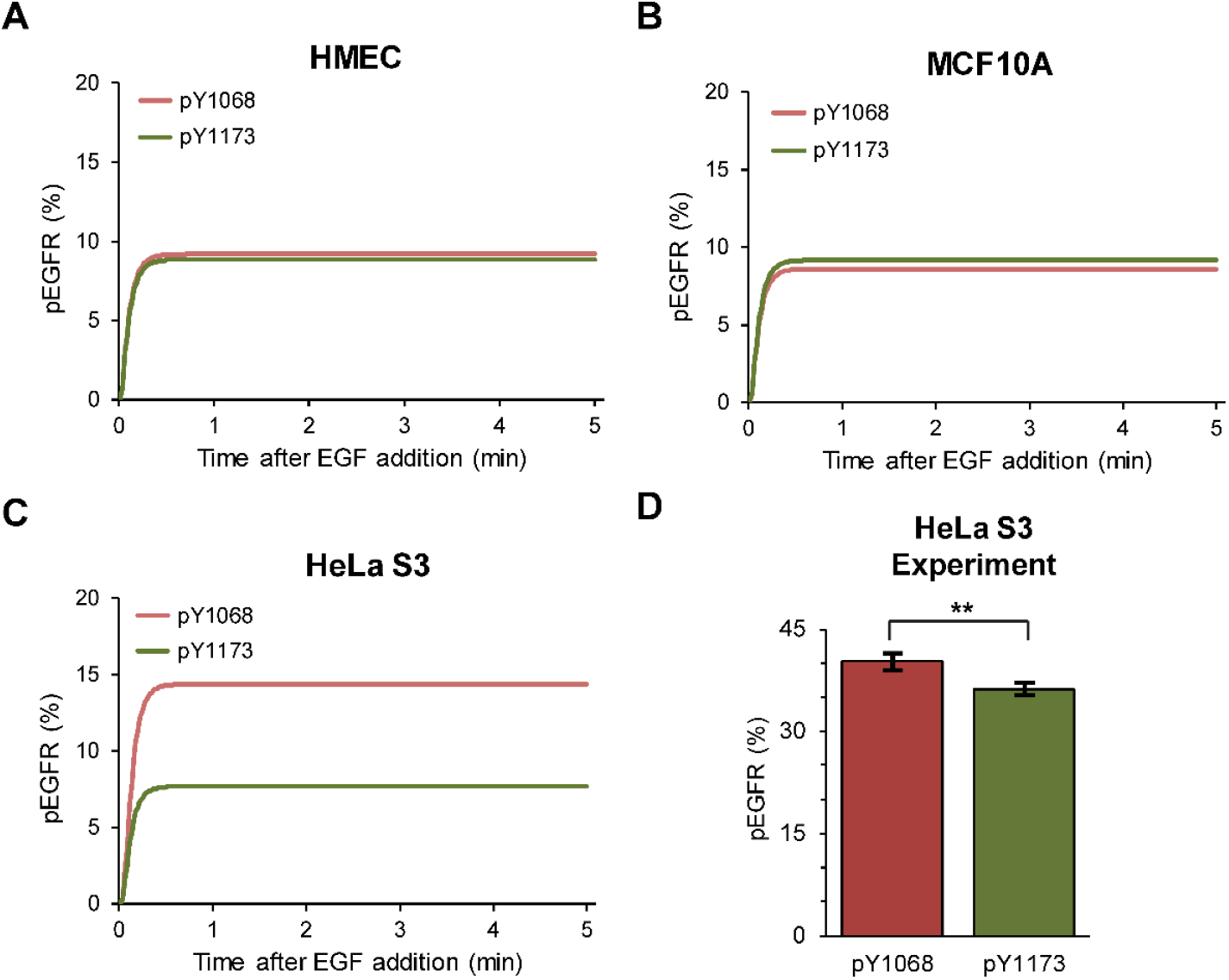
Predicted phosphorylation patterns in cell lines with varying adaptor protein expression. **(A-C)** Predictions of phosphorylation kinetics for MCF10A **(A)**, HMEC **(B)** and HeLa S3 **(C)** cell lines. HMEC cells express low levels of both Grb2 and Shc1; MCF10A cells have slightly higher levels of Shc1 than Grb2; HeLa cells express ∼7 times more Grb2 than Shc1 (Shi et al. 2016; Kulak et al. 2014) (see Supplementary Table S1). **(D)** Phosphorylation pattern in HeLa S3 cells obtained by SiMPull measurements. N>1550. ** P ≤ 0.01. Error bars represent mean +/−S.E.M.

### Three-color SiMPull reveals frequency of multisite phosphorylation on individual EGFR

Our results above suggest that EGF-treated cells bear subpopulations of receptors with differing phosphorylation patterns. SiMPull offers unique advantages over traditional methods, since individual molecules can be probed with more than one antibody providing each has a spectrally distinct fluorescent tag. We utilized the capability for simultaneous three-color SiMPull imaging to determine the frequency of multisite phosphorylation on individual EGFR. Figure 5A illustrates the basic protocol used to evaluate the incidence of multi-site phosphorylation at the single-receptor level. The image in Figure 5B shows a typical result, where each receptor is resolved based upon GFP-emission (Cyan), then overlaid with detection of anti-PY1068-CF555 (green) and detection of anti-PY-AF647 (Magenta). Circled, white spots indicate the presence of EGFR that are positive for both antibodies and, therefore, phosphorylated on PY1068 as well as at least one other tyrosine in the cytoplasmic tail.

**Figure 5.**
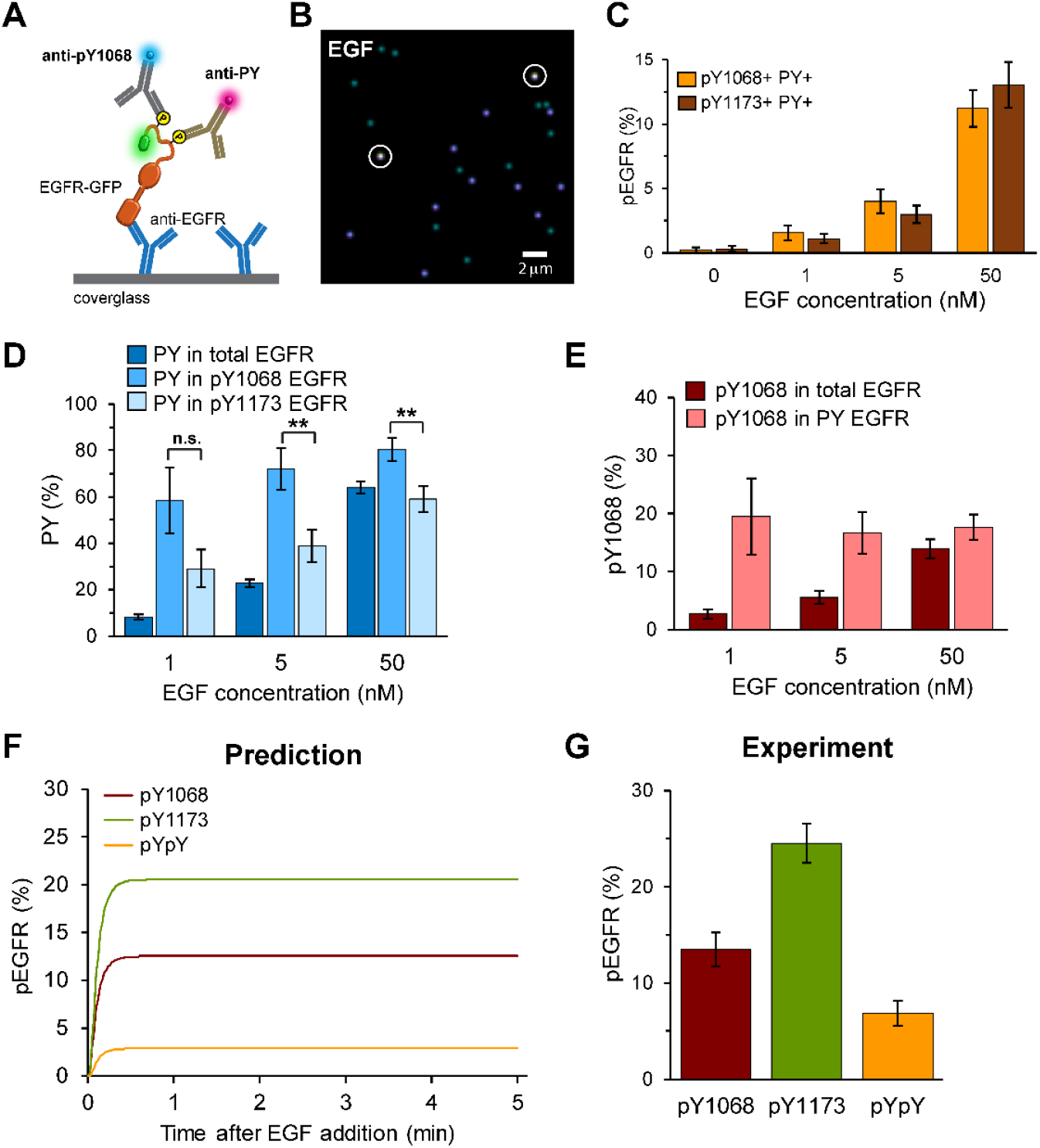
SiMPull reveals EGFR multi-site phosphorylation. **(A)** Schematic of 3-color labeling. **(B)** Representative 3-color SiMPull image of EGFR-GFP (cyan), anti-pan-PY (magenta) and anti-pY1068 (green). White circles indicate receptors that contain all three labels and are therefore positive for both pan-PY and pY1068. This image does not contain receptors labeled with pY1068 alone. Cells were treated with 25 nM EGF for 5 min. **(C)** Quantification of three color SiMPull shows that multi-site phosphorylation is readily observed over a range of EGF concentrations. Three-color images were analyzed for the fraction of EGFR demonstrating co-labeling of pY1068+PY (orange) and for pY1173+PY (red). N>850. **(D)** Plot shows the fraction of pan-phosphorylation found in the total EGFR population (dark blue), the pY1068+ population (medium blue) and the pY1173+ population (light blue). N>850. ** P ≤ 0.01. **(E)** Bar graph showing the fraction of pY1068+ receptors within the total EGFR population (red) as compared to within the phosphorylated receptor population (pink). N>850. **(F)** Mathematical model parameterized by single-site phosphorylation data predicts the presence of dual-site phosphorylation (pYpY, orange line). **(G)** SiMPull experiments demonstrate dual phosphorylation at Y1068 and Y1173 that is consistent with the model prediction. N>750. All error bars represent mean +/−S.E.M.

When labeling a single protein with two or more antibodies, the potential for steric hindrance must be considered. We first tested whether co-labeling of receptors with pairwise combinations of antibodies to PY1068, PY1173 and/or pan-PY was technically feasible. Results in Supplemental Figure S3 confirm the feasibility of a method that employs sequential, pairwise antibody labeling and also provides for correction of steric blocking during our labeling procedures (see also Methods). We also performed step-photobleaching analysis to assure that the doubly labeled receptors were associated with a single EGFR-GFP molecule and not two or more EGFR in a diffraction-limited spot (Supplementary Fig. 4A). Step photobleaching confirmed that at our pull-down density the majority of double-labeled receptors are indeed monomers (∼98%).

By combining anti-pan-PY labeling with each of the two specific sites (pY1173 or pY1068), we were able for the first time to quantify the frequency of multi-site phosphorylation on individual receptors. First, we examined to what extent a receptor phosphorylated at Y1068 or Y1173 might also be phosphorylated at another tyrosine (pan-PY). Figure 5C reports the estimate of the percentage of the total receptor pool exhibiting multi-site phosphorylation, found to reach 13.0 +/−1.8% or higher as a function of dose for each of the two tyrosines examined here. From this data, we can also interrogate the behavior of specific phosphorylation events with respect to a restricted receptor pool. For example, Figure 5D shows the probability of a receptor being phosphorylated (PY+) within three distinct subpopulations. As expected, the total percent of phosphorylated EGFR increased with dose (Fig. 5D, dark blue). At the same time, there is a high probability of a receptor being co-labeled for PY given that is it pY1068+ (P(PY │ pY1068), reaching 80.3 +/−4.9% after 5 min of 50 nM EGF stimulation (Fig. 5D, medium blue). Additionally, we found a high frequency of pY1173+ receptors co-labeled with pan-reactive anti-PY antibodies (Fig. 5D, light blue), albeit lower than for pY1068. These data suggest that, at least in CHO cells, there is a slight increase in multi-site phosphorylation in combination with pY1068 over pY1173. We also analyzed the data in a manner to determine the fraction of pY1068 within the total receptor population (P(pY1068 │ total EGFR)) as compared to the PY+ receptors (P(pY1068 │ PY)). Figure 5E shows that the total fraction of EGFR phosphorylated at Y1068 increases with ligand dose (red), while the fraction of pY1068 within the PY-EGFR population remains relatively constant (pink). Simulations corresponding to experiments in Figures 5D,E are consistent with the observed results (Supplemental Figure 4B,C), and predict that even at sub-nanomolar EGF concentrations the levels of pY1068 in PY-EGFR remain at comparable levels as when EGF concentrations are high (Supplemental Figure 4C). This supports the idea that, at least for tyrosine sites with fast phosphorylation kinetics such as Y1068, the phosphorylation levels within a signaling unit (activated receptor) remain relatively constant and the influence of ligand dose is in the number of activated receptors. Together, these results directly demonstrate that it is common for individual EGFR to acquire phosphorylation on multiple sites.

### Degree of Y1068-Y1173 dual phosphorylation predicted by model is consistent with SiMPull measurements

We next explored the mechanistic basis for multi-site phosphorylation in our computational model, which was initially trained using only single-site phosphorylation data. Simulation results in Figure 5F predict that approximately 3% of receptors will become dually phosphorylated at Y1068 and Y1173 within 1 min of saturating ligand dose, reaching a steady state value sustained for at least 5 min. To test this prediction experimentally, we performed three-color analysis by treating cells for 5 min with EGF, followed by lysis and SiMPull isolation. Samples were co-labeled with anti-pY1068 and anti-pY1173 antibodies and corrected for reduced labeling efficiency due to steric blocking (see Supplementary Fig. 4B-,D and Methods). As shown in Figure 5G, we found that 6.9 +/−1.3% of total receptors have dual phosphorylation at Y1068 and Y1173. As a control, we treated cells with EGF in the presence of phosphatase inhibitors and found a corresponding increase in the percentage of dually-phosphorylated EGFR (Supplemental Fig. S2B). Therefore, both our computational model and quantitative SiMPull experiments support the outcome of dual phosphorylation.

### Dimerization efficiency and lifetime are key factors in multisite phosphorylation

We performed a sensitivity analysis to determine how dual phosphorylation depends on parameter values (Supplementary Fig. S5). Changes in phosphorylation and dephosphorylation rate constants had the greatest effect on the percent of dual phosphorylation. Changes in parameters that govern adaptor protein recruitment had the next largest effects on dual phosphorylation.

To validate the impact of changes in kinase activity, we turned to our prior work with activating EGFR mutations common in non-small cell lung cancer (NSCLC) (Valley et al. 2015). The oncogenic properties of the L858R EGFR mutant have been attributed in part to increased kinase activity (Zhang et al. 2006), as well as structural changes that enhance dimerization potential in the absence of ligand (Valley et al. 2015). Using the mathematical model, we simulated the mono- and dual-site phosphorylation in response to increased kinase activity (reflected by the pseudo first-order rate constant used in the model to characterize autophosphorylation) for L858R EGFR dimers over wild type (WT) EGFR dimers. The model predictions in Figure 6A show that enhanced kinase activity will lead to increases in phosphorylation at both Y1068 and Y1173, as well as in dual phosphorylation. Experimentally, we compared single and multisite phosphorylation in CHO cells expressing WT or L858R EGFR under saturating EGF conditions, where our prior work showed that ligand-bound WT and mutant EGFR have the same dimer lifetime (Valley et al. 2015). Consistent with the modeling predictions, we found that L858R EGFR has an increase in phosphorylation at each tyrosine as well as in Y1068-Y1173 simultaneous phosphorylation (Figure 6B). To obtain a quantitative estimate of the increased kinase activity, we fit our model to the data of Figure 6B. In the fitting procedure, we allowed the rate constant for autophosphorylation to vary, but held all other parameters fixed at their previously estimated values. The one-parameter fit indicates that the intrinsic kinase activity of L858R EGFR is 3.6-fold higher than WT EGFR. Simulation results based on the best fit are shown in Figure 6A. Our estimate is consistent with previous *in vitro* measurements, in which the kinase activity for the L858R kinase domain was approximately 3.5-fold higher (Zhang et al. 2006).

**Figure 6.**
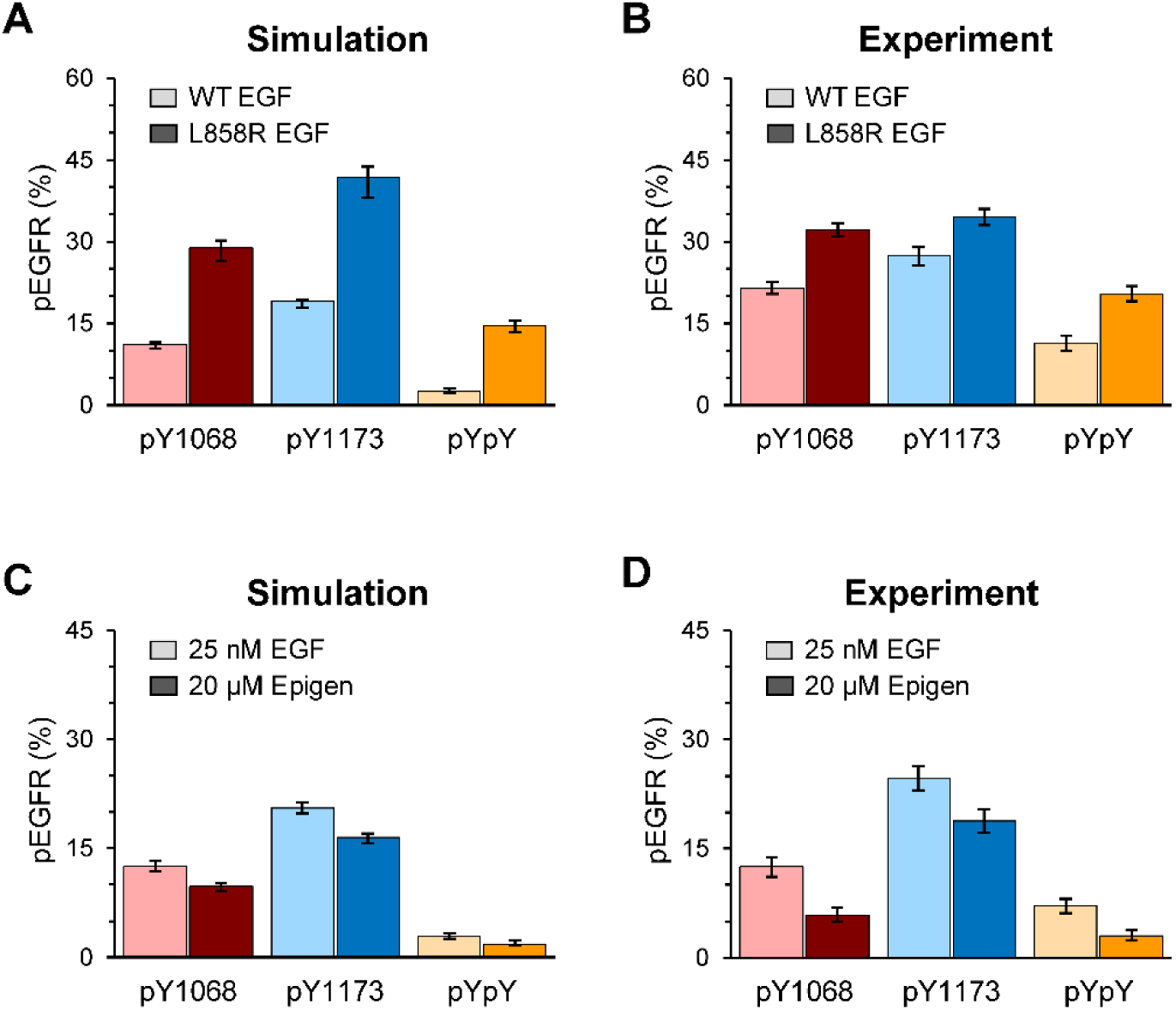
Model predicts altered multi-site phosphorylation resulting from EGFR mutation or low-affinity ligand. **(A)** Model predicts higher single site and multi-site phosphorylation in EGFR-L858R versus WT, where the mutant has a higher kinase activity. **(B)** SiMPull measurements or WT and mutant EGFR expressed in CHO cells confirm the model predictions. Cells were stimulated with 50 nM EGF for 5 min. N>950 **(C)** Model predicts a lower phosphorylation with Epigen stimulation as compared to EGF, where the mechanism is that Epigen induces less stable dimers. **(D)** The model predictions for levels of site-specific and dually phosphorylated EGFR are confirmed by the SiMPull measurements. CHO-EGFR-GFP cells were stimulated for 5 min with the indicated saturating ligand dose. N>1200. All error bars represent mean +/−S.E.M.

We were particularly interested in exploring the interdependent relationships between ligand affinity, dimerization off-rates and multi-site phosphorylation. To explore this experimentally, we focused on comparing multi-site EGFR phosphorylation in cells stimulated with either (high affinity) EGF or another natural EGFR ligand, (low affinity) epigen. We chose this system because shorter dimer lifetimes were recently proposed to be a key feature that explains altered cellular outcomes of epigen-mediated signaling (Freed et al. 2017). We used the mathematical model to predict the outcome of shorter dimer lifetimes on EGFR phosphorylation and found that receptor phosphorylation at pY1068 was reduced (Figure 6C), consistent with qualitative Western Blot results in Freed et al. (2017). The model also predicted that phosphorylation at Y1173 and dual phosphorylation should be reduced. We used 3-color SiMPull to evaluate the differential levels of multi-site phosphorylation for EGFR after stimulation with either EGF or epigen, using saturating levels of both ligands to ensure equivalent occupancy (Figure 6D). We found that both tyrosines were phosphorylated to a lesser extent by epigen than in EGF-stimulated cells. The number of receptors achieving dual-phosphorylation was also reduced with epigen activation. To determine what dimer lifetime best explains these results, we fit our model to the data of Figure 6D. In the fitting procedure, we allowed the rate constant for EGFR dimer dissociation to vary, but held all other parameters fixed at their previously estimated values. The one-parameter fit indicates that epigen-induced dimers have 4.8-fold shorter lifetime than EGF-induced dimers. Simulation results based on the best fit are shown in Figure 6C. These results strengthen the argument that altered epigen signaling is a result of shorter-lived dimer interactions.

## DISCUSSION

Mathematical modeling of cellular signaling, including EGFR signaling, has historically relied on population-based measurements (Kholodenko et al. 1999; B. S. Hendriks et al. 2003). Parameterization of mathematical models has improved over time, incorporating more accurate measurements such as quantitative mass spectrometry techniques to profile overall protein abundance patterns for the EGFR-MAPK pathway (Shi et al. 2015), as well as ligand-induced changes in the phospho-proteome (Yi et al. 2018). Concomitantly, the systems biology field has increasingly recognized the influence of heterogeneity at the single-cell scale in determining signaling output (Kolitz and Lauffenburger 2012). Recent advances in quantitative imaging now provide powerful new methods to interrogate single-cell and single-molecule dynamics, which are providing new insights on how cells encode and interpret information (Welch et al. 2011; Grecco, Schmick, and Bastiaens 2011; Purvis and Lahav 2013). In prior work, we have built spatial stochastic models of EGFR/ErbB signaling (Pryor et al. 2013; McCabe Pryor et al. 2015) that incorporate single-molecule measurements, including experimentally derived values for receptor diffusion and ligand-dependent changes in dimer off-rates (Low-Nam et al. 2011; Steinkamp et al. 2014). These properties of EGFR continue to be key features of our current phosphorylation site-specific, rule-based model.

Wollman and colleagues have proposed that a deeper understanding of signaling networks requires richer data sets collected at the single-cell level, coupled closely with computational modeling in an iterative process of prediction, experimental confirmation and manipulation (Handly, Yao, and Wollman 2016). In this work, we set out to purposely pair simulations with novel single molecule quantitative data, capturing the dose-dependent heterogeneity in phosphorylation of thousands of single receptors. Improvements in the single molecule technique, SiMPull, allowed us to obtain robust, quantitative information about EGFR multi-site phosphorylation patterns and to identify the frequency of subpopulations. These data were used to train a rule-based model of site-specific EGFR phosphorylation and adaptor recruitment. Results of the model were used to guide further experiments in the SiMPull format.

This work has provided several important insights. First, we directly tested the widely-held assumption that SH2 domain binding transiently protects specific phospho-tyrosines from cellular phosphatases, as suggested by prior work using semi-quantitative, population-based western blot techniques (Rotin et al. 1992; Brunati et al. 1998; Sigismund et al. 2013; Jadwin et al. 2018). As our model predicted, overexpression of Grb2 led to a pronounced increase in phosphorylation levels at Y1068 of EGFR, the principal site where Grb2 binds, but spared the Shc1 binding site at Y1173. This enhancement may be particularly relevant in cancer cells, where expression levels of signaling proteins are commonly altered (Santarius et al. 2010; Verbeek et al. 1997). Our combination of modeling and experimentation confirms predictions that modulation of adaptor protein abundance translates into changes in specific receptor phosphorylation patterns (Shi et al. 2016). Although we focused on two prototypical examples of binding partners, Grb2 and Shc1 (Kholodenko, Hoek, and Westerhoff 2000), the principles we describe likely apply to the broader interactome map of ∼ 89 SH2-bearing proteins that interact with ErbB receptors with varying degrees in affinity, abundance and tissue-specific expression (Hause et al. 2012). Of course, some SH2-bearing proteins, such as Grb2, may be somewhat promiscuous and bind with weaker affinity to more than one phospho-site in the receptor cytoplasmic tail (Jadwin et al. 2018). Binding of adaptors at one or more sites may partially block the accessibility of other SH2 proteins or phosphatases at nearby sites or even serve as scaffolds to build larger macromolecular complexes (Hsieh et al. 2010; Telesco et al. 2011). Addressing the combinatorial complexity of these signaling interactions is a strength of rule-based approaches (Chylek et al. 2014), particularly when combined with single molecule quantification.

Second, we demonstrate the advantages of SiMPull for determining the fraction of receptors that bear more than one post-translational modification simultaneously. This ability required considerable optimization of SiMPull protocols, representing a step forward over the SiMBlot approach that failed to detect multisite EGFR phosphorylation (Kim et al. 2016). SiMBlot is based upon surface biotinylation prior to ligand exposure, followed by denaturation and avidin capture for imaging, which may contribute to underestimation of EGF-mediated receptor phosphorylation (Kim et al. 2016). Notably, our modeling and experimental results are consistent with previous work indicating that multisite phosphorylation is important in the efficient recruitment of certain adaptor proteins to activated EGFR (Sigismund et al. 2013; Fortian and Sorkin 2014); it follows that differential phosphorylation patterns connect individual receptors to distinct pathways for biased signaling. It is particularly important to understand biased signaling in the context of different ligand types and doses, as well as cell variables such as the expression levels of receptors and their signaling partners (Chen et al. 2009; Wilson et al. 2012; Freed et al. 2017; Wolf-Yadlin et al. 2006).

Recent work from the Lemmon group has explored differential phosphorylation patterns and signaling outcomes after EGFR stimulation with high and low affinity ligands (Freed et al. 2017). Our contributions to that study supported the conclusion that low affinity ligands, such as epigen, induce distinct dimer structures that are less stable than dimers induced by EGF. By fitting of the rule-based model to SiMPull results derived from epigen-treated cells, we predict that reduced dimer lifetime alone is sufficient to explain the experimentally observed differences in phosphorylation kinetics for the two ligands. Nevertheless, our results here with EGFR mutants indicate that dimer lifetime and kinase efficiency act together to dictate rates of multisite phosphorylation. We showed previously that EGF-bound WT and L858R EGFR dimers have similar lifetimes (Valley et al. 2015), while our new SiMPull results show that EGF treatment leads to significant increases in single- and dual-site phosphorylation for the mutant. Simulations reported in Figure 6 indicate that this can be explained by 3.6-fold increase in intrinsic kinase activity, a result that is reasonably consistent with biochemical measurements using recombinant kinase domains (Zhang et al. 2006). On the other hand, L858R receptors also favor ligand-independent signaling, attributed to “inside-out” signaling that promotes extension of the extracellular domain and exposure of the dimer arm in the absence of ligand (Valley et al. 2015) and/or more favorable orientation of cytoplasmic kinase domains (Shan et al. 2012; Red Brewer et al. 2013). So, the oncogenic properties of the L858R mutation can be attributed to both the increase in catalytic activity (in any pair) and enhanced dimerization in the absence of ligand.

In summary, we have integrated the unique datasets provided by SiMPull with mathematical modeling to better understand the combinatorial aspects of EGFR signaling. Our results provide novel insight into the regulation of EGFR phosphorylation, demonstrating critical roles for adaptor protein abundance and receptor dimerization lifetimes. The abundances of adaptor proteins vary across cell types (Shi et al. 2016). It has been proposed that these variations explain cell-specific functions of EGFR and other RTKs. Here, we have seen that there is an interplay between single-molecule patterns of phosphorylation and adaptor protein abundances. Thus, SiMPull assays complement measurements of these abundances. We have also shown that these types of data can be analyzed in an integrated fashion using a mechanistic model. Further work along these lines will help us to better understand the context-dependent functions of RTKs in the future.

## Acknowledgements

We thank Dr. Donna Arndt-Jovin and Dr. Mark Lemmon for helpful discussions. We also thank Dr. Arndt-Jovin for the CHO-ACP-EGFR cell line and suggesting the ACP-EGFR comparison experiment. We thank Richard Pepermans for generating the Grb2-mCherry construct, Hannah Johnson for generation of the CHO-EGFR-GFP Grb2-mCherry cell line, Shayna Lucero for assistance with cell culture and Aubrey Gibson for assistance with Western Blots. This work was supported by the New Mexico Spatiotemporal Modeling Center (NIH P50GM085273), Mexican Secretariat of Public Education SEP (E.S.-C.), National Science Foundation CAREER MCB-0845062 (D.S.L.), National Institutes of Health R35GM126934 (D.S.L) and the UNM Comprehensive Cancer Center (NIH P30CA118100). We gratefully acknowledge use of the University of New Mexico Comprehensive Cancer Center fluorescence microscopy and flow cytometry facilities, as well as the NIH P30CA118100 support for these cores. We acknowledge use of computing resources provided by the Los Alamos National Laboratory Institutional Computing program, which is operated for the National Nuclear Security Administration of the Department of Energy under contract.

## Author contributions

E.S-C. performed all single molecule experiments and data analysis. E.S-C. and K.A.L. developed analysis methods and algorithms. E.S-C. formulated the mathematical model, estimated model parameter values, and performed simulations and sensitivity analysis with guidance from W.S.H. C.F.N. performed biochemistry experiments. E.D.M. performed bootstrapping and parallel tempering to quantify parameter uncertainty. D.S.L directed the project. D.S.L., E.S-C., K.A.L, and B.S.W. designed and interpreted experiments. D.S.L and E.S-C., W.S.H and B.S.W wrote the manuscript with input from all authors.

## Conflicts of Interest

The authors declare no conflict of interest.

## METHODS

### Cell lines and reagents

CHO cells expressing GFP-tagged(Brock, Hamelers, and Jovin 1999; Lidke et al. 2004) or ACP-tagged EGFR (provided by Dr. Donna Arndt-Jovin) were cultured in DMEM supplemented with 10% FBS, penicillin–streptomycin and 2 mM L-glutamine (Thermo Fisher Scientific). ACP-tagged EGFR was as described in (Valley et al. 2015; Ziomkiewicz et al. 2013) with the exception that a shortened 16 aa sequence was introduced at the EGFR N-terminus(George 2006). EGF, Protease and Phosphatase Inhibitor Cocktail, Alexa Fluor 647 NHS Ester, and NeutrAvidin were purchased from Thermo Fisher Scientific. CoA 488 and ACP Synthase were purchased from New England Biolabs. N-(2-aminoethyl)-3-aminopropyltrimethoxysilane was purchased from United Chemical Technologies (#A0700). Sodium bicarbonate and sodium borohydride were purchased from EMD Millipore (#SX0320-1, #SX0380-3). mPEG-Succinimidyl Valerate (MPEG-SVA-5000-5g) and biotin-PEG-Succinimidyl Valerate (Biotin-PEG-SVA-5000-500mg) were from Laysan Bio. Biotinylated anti-EGFR antibody (E101) was obtained from Leinco Technologies. Antibodies in carrier-free buffer were purchased from Cell Signaling Technologies: EGFR pY1068 (clone 1H12, 2236BF) and EGFR pY1173 (clone 53A5, 4407BF). Monoclonal antibody pre-labeled with AF647 to detect pan-tyrosine phosphorylation (PY99 antibody, sc-7020 AF647) was purchased from Santa Cruz Biotechnology. Mix-n-Stain CF555 and CF640R antibody labeling kits were purchased from Biotium Inc. Paraformaldehyde and glutaraldehyde were purchased from Electron Microscopy Sciences.

### Labeling of antibodies

Carrier-free antibodies (50 μg at 0.5-1 mg/mL per reaction) were labeled using Mix-n-Stain antibody labeling kits following the manufacturer’s instructions. Briefly, the labeling reaction was carried out for 30 min at room temperature and antibodies were centrifuged using the ultrafiltration vial provided to remove the unconjugated dye. Antibodies were resuspended in PBS and stored at 4 °C. The labeling efficiency achieved was between 2.7-4.4 dyes/antibody.

### Cell treatment and lysate preparation

CHO-EGFR-GFP cells were plated overnight in 60 mm tissue culture dishes at 800,000 cells/dish and CHO-ACP-EGFR cells in 24-well plates at 50,000 cells/well. For ACP labeling, CHO-ACP-EGFR cells were washed with serum-free DMEM medium (SFM), incubated with ACP labeling solution (SFM, 10 mM MgCl_2_, 4 μM CoA 488 and 1 μM ACP) for 20 minutes at 37°C and washed three times with SFM previous to stimulation. Cells were washed in Tyrode’s solution (135 mM NaCl, 10 mM KCl, 0.4 mM MgCl_2_, 1 mM CaCl_2_, 10 mM HEPES, 20 mM glucose, 0.1% BSA, pH 7.2) and treated with 25 nM EGF or Tyrode’s solution alone (resting cells) at 37°C. At the indicated time points, cells were placed on ice, washed one time with cold PBS followed by addition of lysis buffer (1% IGEPAL CA-630, 150 mM NaCl, 50 mM Tris pH 7.2) containing Protease and Phosphatase Inhibitors. Cell lysates were collected using cell scrapers (Greiner Bio-One North America, #541070), transferred to fresh tubes on ice and vortexed every 5 min for a total of 20 min. Lysates were centrifuged at 16,000× g for 20 min at 4 °C and the supernatant was transferred to a new tube and stored at −80 °C. For experiments involving treatment of cells with phosphatase inhibitors, cells were pre-treated for 15 min with a Tyrode’s solution containing 1 mM pervanadate (PV) followed by incubation for 5 min in a solution with 50 nM EGF and 1 mM PV. A stock solution of 30 mM PV was prepared before each experiment by mixing equimolar concentrations of hydrogen peroxide and activated sodium orthovanadate that was incubated in the dark for at least 15 min before use.

### Fabrication of hydrophobic arrays and surface functionalization

Coverglasses (24×60mm, #1.5; Electron Microscopy Sciences, #63793) were Piranha-cleaned(Labit et al. 2008) and placed in a coverglass holder (Fisher Scientific, #08-817). Coverglasses were sequentially sonicated in Methanol and Acetone for 10 min each, and in 1M KOH for 20 min using a bath sonicator (Branson Ultrasonics, B1200R-1). These solutions were stored in polypropylene 50 mL tubes (VWR, #89401-564) and reused up to five times. Coverglasses were rinsed with Milli-Q water two times, dried by quickly passing them multiple times over the flame of a Bunsen burner using metal tweezers and placed in a dry coverglass holder. A solution containing 76 mL of methanol, 4 mL of acetic acid and 0.8 mL of aminosilane (N-(2-aminoethyl)-3-aminopropyltrimethoxysilane) was prepared in an Erlenmeyer flask, immediately poured into the coverglass holder and incubated at room temperature for 10 min in the dark, followed by 2 min sonication and another 10 min incubation in the dark. Coverglasses were next washed with methanol for 2 min, rinsed and washed for 2 min with water, and dried in the dark. Treated coverglasses were placed on top of a parafilm-covered coverglass containing a guide pattern, which was used as reference to draw the Sample Array with a hydrophobic barrier pen (Vector Laboratories, #H-4000). Ink was allowed to dry for at least 5 min before coverglasses were placed in a humidified chamber (empty tip rack with 50 mL of water; USA Scientific #1111-2820). For surface functionalization, 50 mg of mPEG-Succinimidyl Valerate, 1.3 mg of biotin-PEG-Succinimidyl Valerate and 200 µL of freshly prepared 10 mM sodium bicarbonate were mixed thoroughly by pipetting up and down, centrifuged for 1 min at 10,000 g at room temperature and immediately applied to the SiMPull array (10-13 uL per region). After incubating for 3-4 hours in the dark inside the humidified boxes, arrays were washed by sequential 30 sec submersions into three water-filled 250 mL glass beakers. Coverglasses were dried with nitrogen gas, stored in pairs (back to back) inside 50 mL tubes, which were filled with nitrogen gas before closing and sealing with Parafilm. Coverglasses were stored in the dark at −20°C for up to a week before use.

### Labeling and quantification of surface receptors

CHO-EGFR-GFP cells grown in 24-well plates were placed on ice and washed 3 times with cold PBS. AF647-NHS Ester was dissolved at the indicated concentrations in PBS. Cells were incubated with this solution for 30 min at 4°C with gentle agitation, washed 3 times with cold PBS and subjected to cell lysis. The percent of receptors labeled with AF647 across different dye concentrations was assessed with SiMPull. To estimate the percent of receptors at the cell surface the AF647-labeling curve was fitted to a biexponential decay curve in its increasing form using the ‘fit’ function in MATLAB: y = C1 (1 - e^-ax^) + C2 (1 - e^-bx^), where *y* is the % of AF647-labeled receptors, *x* is the concentration of reactive AF647-NHS ester used, and *a*>0, *b*>0, *C1* and *C2* are coefficients to be fitted. The sum of the coefficients *C1* and *C2* represent the asymptote of the curve and an approximation of the fraction of receptors at the cell surface.

### Single-Molecule Pulldown and phospho-site labeling

T50 (10 mM Tris pH 8.0, 50 mM NaCl) and T50-BSA (T50 with 0.1 mg/mL BSA) solutions were prepared and stored for up to a month at room temperature. SiMPull arrays were equilibrated at room temperature and placed on a TC100 plate lined with Parafilm. Each region of the SiMPull array was treated with 10-15 µL of a 10 mg/mL sodium borohydride (NaBH_4_)/PBS solution for 4 min at room temperature and washed 3 times with PBS. SiMPull regions were then incubated with a 0.2 mg/mL NeutrAvidin/T50 solution for 5 min and washed three times with T50, followed by incubation with a 2 µg/mL biotinylated anti-EGFR/T50-BSA solution for 10 min and washed three times with T50-BSA.

The plate containing the SiMPull array(s) was kept on ice during sample preparation. Lysates were diluted in cold T50-BSA with Protease and Phosphatase Inhibitors (T50-BSA/PPI), vortexed at medium speed, and added to the SiMPull array. After 10 min incubation, the lysates were removed and the SiMPull regions washed 4 times with cold T50-BSA/PPI. To determine appropriate dilution factor, the density of pulldown receptors as a function of lysate concentration was first assessed to achieve a pulldown density 0.04-0.08/μm_2_. Antibodies were diluted in cold T50-BSA/PPI, incubated for 1 hr, washed 6 times with cold T50-BSA for a total of 6-8 minutes, and washed twice with cold PBS. Immediately after, antibodies were fixed for 10 min with a 4% PFA/0.1% GA solution (paraformaldehyde/glutaraldehyde) and washed 2 times with 10 mM Tris (pH 7.4)/PBS for a total of 10 min to inactivate fixatives. For 3-color SiMPull experiments the same antibody incubation and fixation procedure was performed for the second antibody. Tris solution was replaced by T50-BSA and the SiMPull array was equilibrated to room temperature before proceeding to imaging. The antibodies used for detection of EGFR pY1068 (clone 1H12) and pY1173 (clone 53A5) have a high specificity for their target sites (Kim et al. 2016).

### SiMPull imaging

Imaging of SiMPull samples was performed using an inverted microscope (Olympus America, model IX71) equipped with a 150×/1.45 NA oil-immersion objective for Total Internal Reflection Fluorescence Microscopy (Olympus America, UAPON 150XOTIRF) and a three-dimensional piezostage (Mad City Labs, Nano-LPS100). Excitation of CF640R- or AF647-labeled antibodies was done using a 642-nm laser (Thorlabs, HL63133DG), CF555-labeled antibodies using a 561-nm laser (Coherent Inc, Sapphire 561-100 CW CDRH), and of GFP- and CoA 488-tagged receptors using a 488-nm laser (Spectra Physics, Cyan 100mW). All lasers were set in total internal reflection configuration, and laser powers were adjusted to prevent photobleaching of the sample at the timescale of the image exposure time (300 msec). Sample illumination and emission were filtered using a quad-band dichroic and emission filter set (Semrock, LF405/488/561/635-A-000). Emission light was separated into four channels using a quad-view multichannel imaging system (Photometrics, model QV2) equipped with the appropriate dichroics (Chroma, 495 DCLP, 565 DCLP, 660 DCLP) and emission filters (Semrock, 685/40 nm, 600/37 nm, 525/45 nm). Emission light was collected with an electron-multiplying charge-coupled device (EMCCD) camera (Andor Technology, DU-897E-C50-#BV) with EM gain set to 200. Each channel was 256 x 256 pixels, with a pixel size of 106.7 nm. Photobleaching and bleed through were prevented by controlling the laser shutters and microscope stage through a MATLAB script to sequentially excite and acquire the different fluorophores (642-nm laser first, 488-nm laser last). A minimum of 20 regions of interest were acquired per condition. For quantification of step photobleaching of EGFR-GFP molecules, a 100 frame time series (300 msec exposure time) was acquired after imaging of the other two channels.

### Quantification of Receptor Phosphorylation

All image processing was performed using MATLAB together with the MATLAB toolbox for image-processing DIPImage (Delft University of Technology)(C. L. L. Hendriks et al. 1999) and all software is available upon request. The location of emitters in each channel was calculated using graphics processor unit (GPU) computing as previously described(Smith et al. 2010). Fits in the GFP channel were filtered based on the quality of the fit to the point spread function to reduce the chances of detecting multiple receptors in close proximity as a single molecule. Image registration was performed as previously described (Schwartz et al. 2017). In our case, the root mean square error for image registration was <10 nm. For visualization purposes, Gaussian blob representations of the fluorophore localizations were generated. A receptor was considered to be phosphorylated when the localization centers of the receptor and labeled antibody were at a distance <106.7 nm (within 1 pixel).

Phosphorylation percentages were calculated as 100*(*N*_Phos_)/(*N*_GFP_-*N*_BG_) where *N*_Phos_ is the number of receptors identified as phosphorylated, *N*_GFP_ is the number of observed single molecules in the GFP channel and *N*_BG_ is the non-specific background rate in the GFP channel.

The number of GFP localizations was calculated by subtracting background spots and accounting only for surface receptors as follows: *N*_GFP_ = (*N*_LOC_-*N*_BG_)\**SR*, where *N*_LOC_ is the total number of emitters localized, *N*_BG_ is the expected number of background emitters in the area imaged, and *SR* (surface ratio) is the fraction of receptors located at the cell surface. The density of background emitters was quantified for each SiMPull array and used for background correction of samples in that array. For 3-color SiMPull experiments where steric hindrance between sequentially incubated antibodies was observed (i.e. pY1068-pY1173 detection), estimations of dual phosphorylation were corrected to account for this hindrance as explained in Supplementary Figure 3. Note that use of a three-color imaging scheme to correlate phospho-antibody labeling directly with GFP-tagged receptors was critical, due to the relatively high non-specific binding of the antibodies (Supplementary Fig. S6). In the absence of the GFP channel to remove the non-specific binding, the values for dual labeling are underestimated (Supplementary Fig. S6).

### Statistical Analysis

Based on the consideration that the phosphorylation state of each receptor analyzed has the properties of a Bernoulli trial, standard errors (SE) of phosphorylation measurements were calculated as for sample proportions in a binomial distribution: *SE*= *p*(1-*p*)/*n*, where *p* is the fraction of receptors phosphorylated and *n* is the total number of receptors. The condition np>10 (with the exception of Figure 6E, np>5) and np(1-p)>10 was ensured to be met to allow this approximation to be adequate. Two-sample Z-test (two-tailed) was used to estimate p-values(LeBlanc 2004).

### Step-photobleaching Analysis

For step-photobleaching analysis of multi-phosphorylated receptors the average fluorescence intensity of the area (200×200 nm) surrounding each of these EGFR-GFP molecules was quantified and plotted for the duration of the time series. Intensity plots were manually analyzed and the number of photobleaching steps was quantified. For a small fraction of the emitters the number of molecules could not be reliably counted because either they photobleached too quickly (<2 frames) or did not photobleach during the duration of the movie, and therefore were excluded from the analysis.

### Mathematical Modeling

The model was formulated using the BioNetGen language (BNGL) (Faeder, Blinov, and Hlavacek 2009). The BioNetGen input file that defines the model is available as Supplementary File S1. This file is a plain-text file. The model accounts for the proteins, sites, and interactions depicted in Figure 2B. The model implicitly accounts for three compartments, which are each taken to be well-mixed: the extracellular fluid, the plasma membrane, and the cytosol. In the model, EGFR dimer-dependent autophosphorylation is taken to be asymmetric, as discussed in the main text. In addition to Y1068 and Y1173, the model accounts for other tyrosines in EGFR, which are lumped together and labeled YN in the BNGL-formatted model-definition file. The model structure was optimized using model restructuration (reviewed in the supplementary tutorial of (Erickson et al. 2019).

In the one-parameter fits carried out to characterize the increased kinase activity of L858R EGFR and the lifetime of epigen-induced EGFR dimers, we used simple grid search as the fitting method, meaning that we systematically varied the free parameter over a specified range, calculating the quality of fit at each parameter value. When the free parameter was the rate constant for autophosphorylation, we considered a range 0.1x below and 100x above the estimated value for WT EGFR. We considered a step size of 0.1 in the grid search. When the free parameter was the rate constant for EGFR dimer dissociation, we similarly considered a range 0.1x below and 100x above the estimated value for EGF-induced dimers. We again considered a step size of 0.1 in the grid search.

Simulations were performed as follows. In a preprocessing step, rules were interpreted by BioNetGen to obtain the rule-implied reaction network and the corresponding ordinary differential equations (ODEs) for mass-action chemical kinetics. Simulations of the dynamics of responses to ligand stimulation were performed by numerically integrating the BioNetGen-derived ODEs using CVODE with BioNetGen’s default CVODE settings. CVODE (https://computation.llnl.gov/projects/sundials) (Hindmarsh et al. 2005) is a dependency of BioNetGen.

Fitting was performed using the differential evolution (DE) algorithm implemented in PyBioNetFit version 0.3.2 (https://github.com/lanl/PyBNF) (Mitra et al. 2019). An important parameter of the DE algorithm is population size: this parameter was set to 200. Fitting runs were allowed to continue until apparent convergence, i.e., until the value of the objective function in the optimization problem stopped decreasing. PyBioNetFit (Mitra et al. 2019) is a software package that replaces BioNetFit (Thomas et al. 2016). PyBioNetFit takes as input three file types: a BioNetGen input file (i.e., a BNGL-formatted model definition), a configuration file, and one or more EXP files with experimental data. These files are packaged together in a ZIP archive as Supplementary File S2. The parameters listed in Supplementary Table S2 were taken to be the free parameters to be adjusted by PyBioNetFit during fitting within the indicated intervals. All other parameters were held fixed at their estimated values.

Confidence intervals for best-fit parameter values were estimated using PyBioNetFit’s bootstrapping procedure. A total of 1000 bootstrap replicates of fitting were performed, and the reported 90% confidence intervals give the range from the 5_th_ percentile to the 95_th_ percentile of the replicate results (Supplementary Table S3). The bootstrapping procedure used the same BioNetGen and EXP files as before and the configuration file fit_bootstrap.conf provided in Supplementary File S2.

Bayesian uncertainty quantification (UQ) was performed using the parallel tempering (PT) algorithm implemented in PyBioNetFit, which is a parallelized Markov chain Monte Carlo (MCMC) method. We used PT to find 90% credible intervals for each parameter listed in Supplementary Table S2. Ten independent runs were performed, each consisting of 200,000 iterations (36 simulations per iteration), with samples recorded after a burn-in period of 50,000 iterations. Reported 90% credible intervals (Supplementary Table S3) for each parameter give the range from the 5_th_ to 95_th_ percentile of the sampled parameter sets (Note that credible intervals are not identical to bootstrap confidence intervals). Prediction uncertainties (Fig. 2D, shaded areas; Fig. 6A and 6C, error bars) were calculated by running simulations with sampled parameter sets and finding the 2.5th and 97.5_th_ percentile pEGFR percentage at each time point. UQ jobs used the same BioNetGen and EXP files as before and the configuration file fit_pt.conf provided in Supplementary File S2. Distributions of parameter values in sampled parameter sets and the correlation between each pair of parameters (Supplementary Fig. S7) were plotted using the function plotmatrix from MATLAB.

Fitting was performed on a laptop. Bootstrapping was performed using 7,000 core hours on a multi-core workstation. Parallel tempering was performed using 40,000 core hours on a computer cluster.

## Notes

#### Summary of Updates

This version includes additional experimental results and mathematical modeling exploring multisite phosphorylation in EGFR, and the impact of oncogenic mutations and ligands with varying affinity on phosphorylation kinetics.

https://github.com/RuleWorld/RuleHub/tree/2019Jun18/Published/Salazar-Cavazos2019

