## Supplementary Tables and Figures for "Integrating Multiplex SiMPull and Computational Modeling to Evaluate Combinatorial Aspects of EGFR Signaling"

### SUPPLEMENTARY MATERIAL

#### Supplementary Table S1. Fixed parameters (not allowed to vary in fitting).

| Parameter description | Name in model | Value | Reference |
|---|---|---|---|
| Avogadro constant | NA | 6.02214 x $10^{23}$ molecules / mol | -- |
| Cytoplasmic volume of mammalian cell (HeLa) | Vc | 1.0 x $10^{-12}$ L | (Fujioka et al. 2006) |
| Extracellular volume per cell (in vitro) | Vextra | 1.0 x $10^{-9}$ L | This study |
| EGF concentration in extracellular media | EGFconc | 0-50.0 x $10^{-9}$ M | This study - particular to each condition |
| Association rate constant for adaptor protein binding to pY sites in EGFR [a] | kp ($k^+$) | 5.0 x $10^6$ $M^{-1}$ $s^{-1}$ | Consistent with (Morimatsu et al. 2007) |
| Association rate constant for EGF-EGFR interaction [a] | kp_EGF | 8.0 x $10^6$ $M^{-1}$ $s^{-1}$ | Set to resemble EGFR phosphorylation kinetics observed in (Reddy et al. 2016) |
| Equilibrium dissociation constant for EGF-EGFR interaction [a, b] | Kd_EGF | 2.0 x $10^{-9}$ M | Consistent with (Björkelund, Gedda, and Andersson 2011) |
| Rate constant for dissociation of EGFR dimers (each bound to an EGF molecule) | km_dim_L_L | 0.273 $s^{-1}$ | (Low-Nam et al. 2011) |
| Equilibrium dissociation constant for EGFR-EGFR interaction [c] | KD_dim | EGFR_total / 20 | Set so EGFR_total / KD_dim >> 1 |
| Equilibrium dissociation constant for Grb2 SH2 domain-pY1068 EGFR interaction [a, c] | Kd_GE | 0.6 x $10^{-6}$ M | (Morimatsu et al. 2007) |
| Equilibrium dissociation constant for Shc1 PTB domain-pY1173 EGFR interaction [a, c] | Kd_SE | 0.6 x $10^{-6}$ M | Assumed to be the identical as for Grb2, based on (Hause et al. 2012) |
| Relative rate of phosphorylation of pY sites in receiver receptor vs. activator receptor. | -- | 0.7 (dimensionless ratio) | Estimated from (Kovacs et al. 2015) |

**Supplementary Table S1 (continued)**

| Parameter description | Name in model | Value | Reference |
|---|---|---|---|
| EGFR abundance in **CHO** cells | EGFR_total | $7.7 \times 10^5$ molecules / cell | This study |
| EGFR abundance in **HMEC** cells | EGFR_total | $3.54 \times 10^5$ molecules / cell | (Shi et al. 2016) |
| Grb2 abundance in **HMEC** cells | GRB2_total | $0.43 \times 10^5$ molecules / cell | (Shi et al. 2016) |
| Shc1 abundance in **HMEC** cells | SHC1_total | $0.25 \times 10^5$ molecules / cell | (Shi et al. 2016) |
| EGFR abundance in **MCF10A** cells | EGFR_total | $2.29 \times 10^5$ molecules / cell | (Shi et al. 2016) |
| Grb2 abundance in **MCF10A** cells | GRB2_total | $0.50 \times 10^5$ molecules / cell | (Shi et al. 2016) |
| Shc1 abundance in **MCF10A** cells | SHC1_total | $0.81 \times 10^5$ molecules / cell | (Shi et al. 2016) |
| EGFR abundance in **HeLa** cells | EGFR_total | $0.93 \times 10^5$ molecules / cell | (Kulak et al., 2014) |
| Grb2 abundance in **HeLa** cells | GRB2_total | $6.28 \times 10^5$ molecules / cell | (Kulak et al., 2014) |
| Shc1 abundance in **HeLa** cells | SHC1_total | $1.12 \times 10^5$ molecules / cell | (Kulak et al., 2014) |

[a] Concentration was converted from molar units to (molecules / cell).

[b] Dissociation rate constant was estimated by dividing the equilibrium dissociation constant by the association rate constant ($k^- = K_D / k^+$).

[c] Association rate constant was estimated dividing the dissociation rate constant by the equilibrium dissociation constant ($k^+ = k^- / K_D$).

**Supplementary Table S2. Free (adjustable) parameters.**

| Parameter description | Name in model | Optimization range | Best-fit value [a] | Reference for feasible range |
|---|---|---|---|---|
| Grb2 abundance | GRB2_total | $1.0 \times 10^4$ - $1.0 \times 10^6$ molecules / cell | $1.70 \times 10^5$ molecules / cell | (Shi et al. 2016; Kulak et al. 2014) |
| Shc1 abundance | SHC1_total | $1.0 \times 10^4$ - $1.0 \times 10^6$ molecules / cell | $6.49 \times 10^5$ molecules / cell | (Shi et al. 2016; Kulak et al. 2014) |
| Rate constant for dephosphorylation at sites Y1068 and Y1173 | kdephosY1068 and kdephosY1173 | 0.1 - 100.0 $s^{-1}$ | 1.66 $s^{-1}$ | (Kleiman et al. 2011) |
| Rate constant for phosphorylation : rate constant for dephosphorylation (for Y1068 and Y1173) | ratio_kpkd_Y1068 and ratio_kpkd_Y1173 | 0.01 - 100.0 | 0.158 | (Kim et al. 2012; Kleiman et al. 2011) |
| Rate constant for dephosphorylation at sites other than Y1068 and Y1173 (i.e., at sites lumped together and labeled 'YN') | kdephosYN | 0.001 - 100.0 $s^{-1}$ | 0.017 $s^{-1}$ | (Reddy et al. 2016; Kleiman et al. 2011) |
| Rate constant for phosphorylation : rate constant for dephosphorylation (for 'YN') | ratio_kpkdYN | 0.01 - 100.0 | 0.445 | (Kim et al. 2012) |

[a] Best-fit was found using the synchronous differential evolution (DE) optimization algorithm implemented in PyBioNetFit.

**Supplementary Table S3. Confidence/credible intervals on parameter estimates.**

| Name in model | 90% confidence intervals from bootstrapping procedure | 90% credible intervals from parallel tempering |
|---|---|---|
| GRB2_total | $1.02 \times 10^4$ - $3.90 \times 10^5$ molecules / cell | $1.18 \times 10^4$ - $3.11 \times 10^5$ molecules / cell |
| SHC1_total | $2.08 \times 10^5$ – $1.00 \times 10^6$ molecules / cell | $3.65 \times 10^4$ – $9.06 \times 10^5$ molecules / cell |
| kdephosY1068 and kdephosY1173 | 0.10 - 95.17 $s^{-1}$ | 0.30 – 68.88 $s^{-1}$ |
| ratio_kpkd_Y1068 and ratio_kpkd_Y1173 | 0.113 - 0.278 | 0.126 – 0.221 |
| kdephosYN | 0.004 - 62.58 $s^{-1}$ | 0.015 – 0.020 $s^{-1}$ |
| ratio_kphosYN | 0.314 - 0.762 | 0.422 – 0.471 |

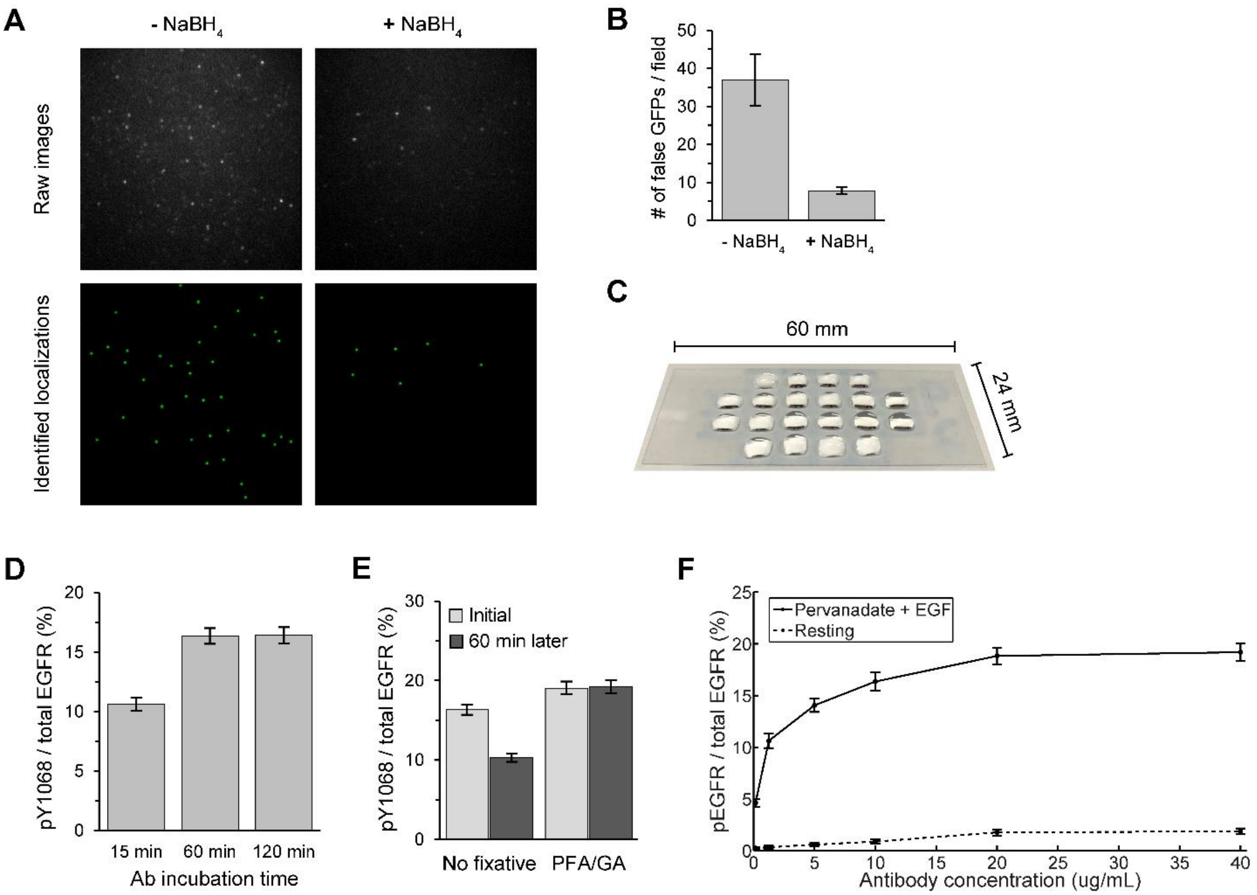


**Supplemental Figure S1. SiMPull Optimization. (A-B)** Autofluorescence is reduced with Sodium Borohydride ($NaBH_4$) treatment. **(A)** Raw images and blob-reconstructions from a typical field of view of a PEG/PEG-biotin functionalized surface without (left) and with (right) $NaBH_4$-treatment. **(B)** Quantification of the average number of false-positive localizations per field of view in surfaces with or without treatment with $NaBH_4$. For each condition N > 12 fields of view were analyzed. Error bars represent mean +/- S.E.M. **(C)** Hydrophobic array for preparation of SiMPull samples. **(D-F)** CHO-EGFR-GFP cells were pre-treated with 1 mM PV for 15 min and stimulated with 50 nM EGF+1mM PV for 5 min at 37°C to enhance receptor phosphorylation and interrogated for anti-EGFR-pY1068-CF555 labeling. **(D)** Antibody labeling with anti-pY1068 requires 60 min to reach maximal labeling. A 20 μg/mL antibody concentration was used. Number of receptors analyzed per condition, N>3400. **(E)** Addition of PFA/GA post-fixation prevents loss of antibody over time. N>2700 per condition. **(F)** Increase in labeling as a function of antibody dose. EGFR-pY1068-CF555 saturates at ~20 μg/mL. Antibody was incubated for 1 hour on ice and post-fixed with PFA/GA. Resting cells were used as a control for non-specific labeling. N>1700 per data point. All error bars are standard error of measured phosphorylation percentages.

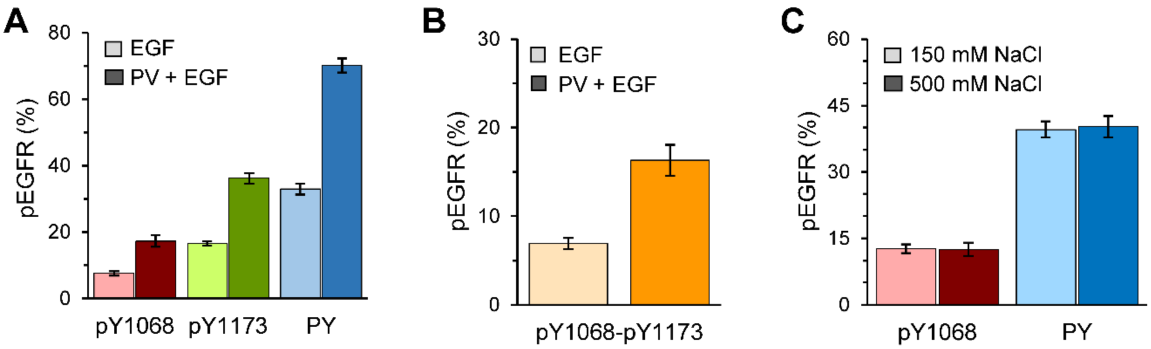


**Supplementary Figure S2. Effect of phosphatase inhibition or cell lysate salt concentration on detected phosphorylation levels. (A)** CHO-EGFR-GFP cells were stimulated at 37°C with either 50 nM EGF for 5 min or pre-treated with 1 mM pervanadate (PV) for 15 min and then stimulated with 50 nM EGF and 1mM PV (PV + EGF) for 5 min. Considering that pervanadate treatment induces EGFR phosphorylation that may not be restricted to the plasma membrane, no surface correction was applied for this figure. PV treatment increases the fraction of phosphorylated EGFR detected by each antibody. Number of receptors per condition, 690 < N < 3400. **(B)** Dual-site phosphorylation is also increased with pervanadate treatment. **(C)** CHO-EGFR-GFP cells were stimulated at 37°C with 25 nM EGF for 1 min and protein extraction was performed with either regular lysis buffer containing 150 mM NaCl (see Methods) or 500 mM NaCl. High NaCl concentrations have been shown to promote disruption of interactions between SH2-containing proteins and their phosphorylated binding partner sites (Grucza et al. 2000). 670 < N < 1600. Error bars are standard error of measured phosphorylation percentages.

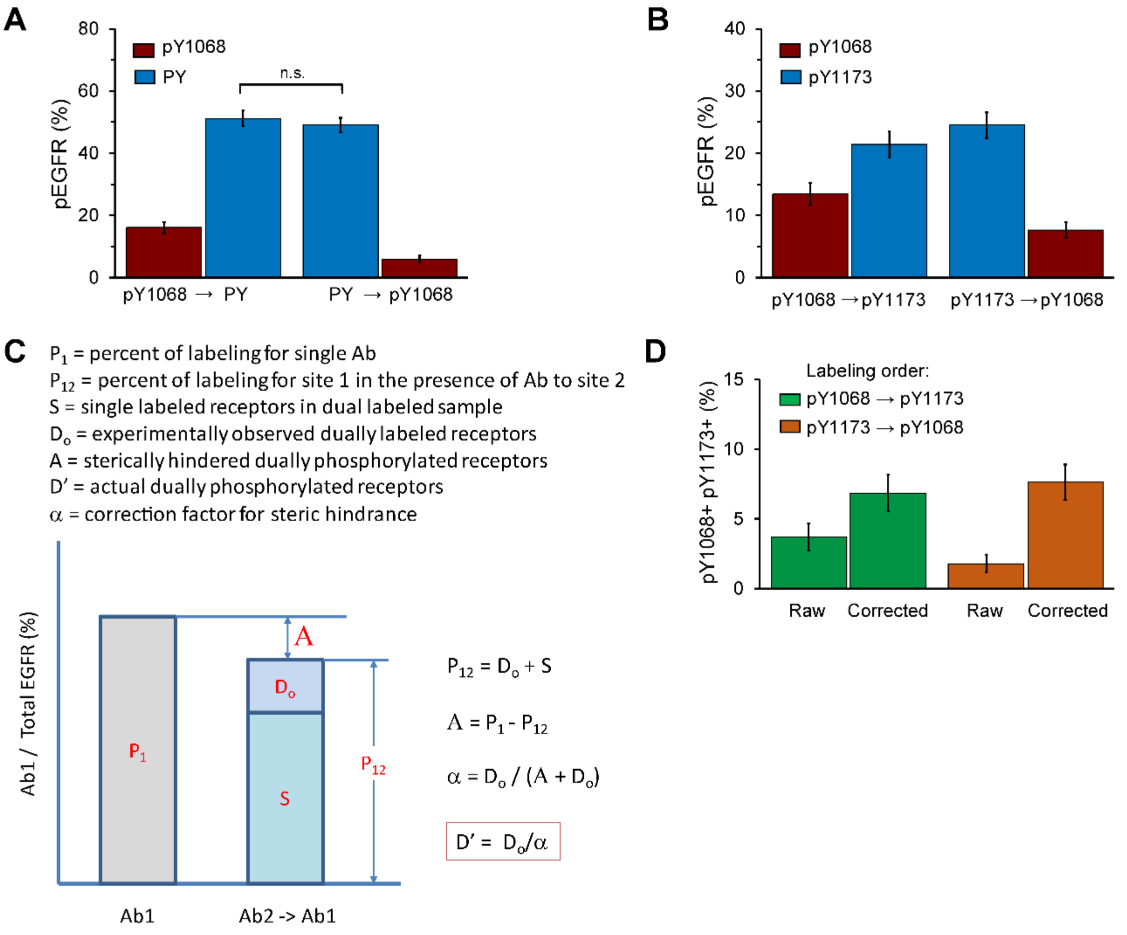


**Supplementary Figure S3. Assessment and correction of steric hindrance in sequentially incubated antibodies for 3-color SiMPull. (A)** Evaluation of steric hindrance between anti-pY1068-CF555 and anti-PY-AF647 (PY) antibodies. CHO-EGFR-GFP cells were stimulated with 25 nM EGF for 5 min at 37°C and EGFR phosphorylation quantified using 3-color SiMPull. Labeling with anti-pY1068 first did not reduce subsequent labeling by anti-PY (pY1068 → PY). However, a reduction in pY1068+ receptors is seen when the labeling order is reversed (PY → pY1068). Number of receptors analyzed per measurement, N>800. n.s. not significant, P = 0.5187. **(B)** Evaluation of steric hindrance between anti-pY1068-CF555 and anti-pY1173-CF640R antibodies. Cells were stimulated as described in (A) and receptor phosphorylation assayed by 3-color SiMPull. In this case, a reduction in labeling was observed for the antibody that is applied second in the labeling sequence. N>780 per measurement. **(C)** Diagram describing estimation of correction factor (*α*) to calculate actual fraction of receptors with dual phosphorylation (*D'*). The observed reduction in labeling with Antibody 1 alone (left bar) as compared to Antibody 1 following Antibody 2 (right bar) indicates the level of steric hindrance. From this information, the correction factor can be calculated. **(D)** Validation of the correction factor by exchanged labeling order. After applying the correction factor ("Corrected" bars), the percentage of pY1068+pY1173+ receptors is similar between experiments where labeling was reversed. All error bars are standard error of measured phosphorylation percentages.

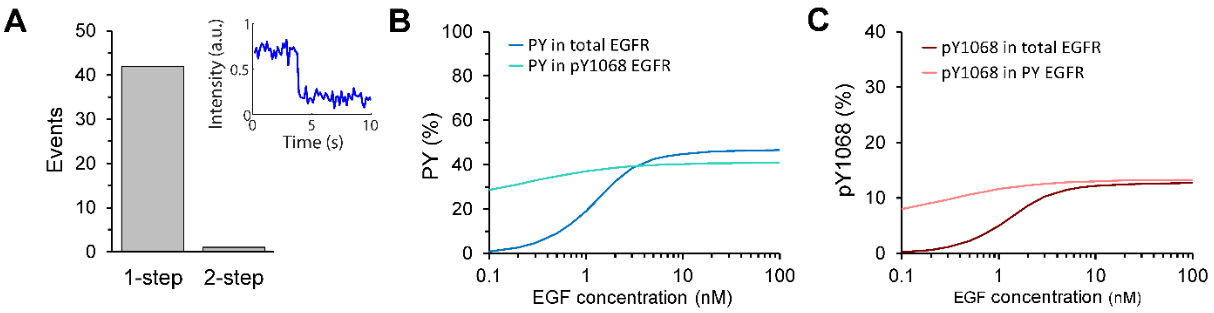


**Supplementary Figure S4. Observed multisite phosphorylation is not an artifact of EGFR dimers and simulations are consistent with experimental observations. (A)** Step-photobleaching analysis of multi-phosphorylated EGFR-GFP. The majority (98%) of diffraction limited GFP spots show single-step bleaching, consistent with the pull-down of receptors as monomers. Inset shows example GFP-intensity trace of a multi-phosphorylated EGFR-GFP demonstrating a single GFP photobleaching step. It is important to note that the number of GFP spots demonstrating two-step photobleaching increased as the sample density increased (data not shown). Therefore, we recommend a pulldown protein density in the range of 0.04-0.08/μm$^2$. Alternatively, photobleaching traces can be performed in each measurement to exclude those spots showing more than one-step photobleaching. **(B,C)** Simulations corresponding to Figures 5D,E. As can be seen in (C), the model predicts that the percent of pY1068 in PY EGFR (labeled with pan-PY antibody) is insensitive to EGF dose, consistent with Figure 5E.

GRB2, EGFR and SHC1 = protein copy numbers per cell
kp_EGF and km_EGF = EGF ligand association and dissocation rate constants
kp_dim and km_dim = rate constants for EGFR dimerizarion and dimer breakup
kp_GE and km_GE = association and dissociation rate constants for Grb2-pEGFR complex
kp_SE and km_SE = association and dissociation rate constants for Shc1-pEGFR complex
kdephos and kphos = rate constants for dephosphorylation and phosphorylation of tyrosine residues in EGFR

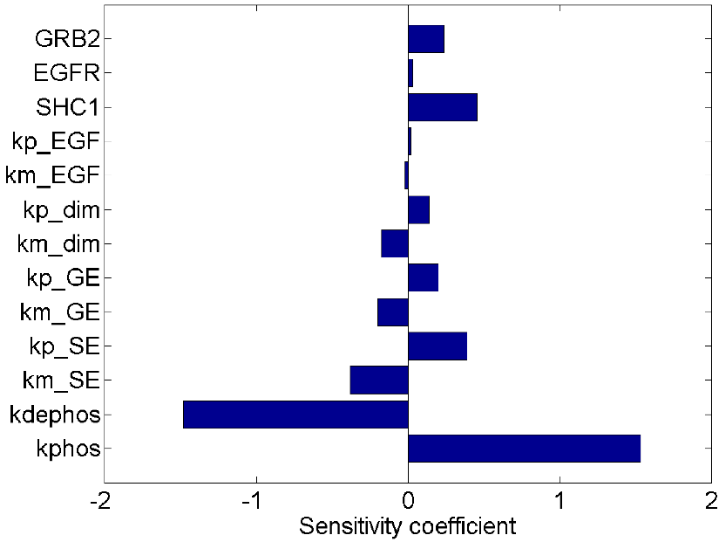


**Supplementary Figure S5. Parameter sensitivity analysis to assess influence of each parameter on multisite phosphorylation.** Simulations were performed to assess the impact on model output, when varying key parameters such as adaptor expression levels (Grb2, Shc1) and their on/off rate constants (kp_GE; km_GE; kp_SE; km_SE), rate constants for ligand association/dissociation (kp_EGF; km_EGF), rate constants for dimerization and dimer breakup (kp_dimer; km_dimer), and rate constants for phosphorylation/dephosphorylation (kphos; kdephos). In this analysis, we increased the nominal value of each parameter in the model by a small amount (1%) and calculated the new level of dual phosphorylation. The results from this analysis were used to calculate sensitivity coefficients, each of which is defined as *SC*= *(∆pYpY/pYpY)/(∆parVal/parVal);* where *pYpY* is the value for nominal parameter values, ∆*pYpY* is the change and *(∆parVal/parVal) =0.01.*

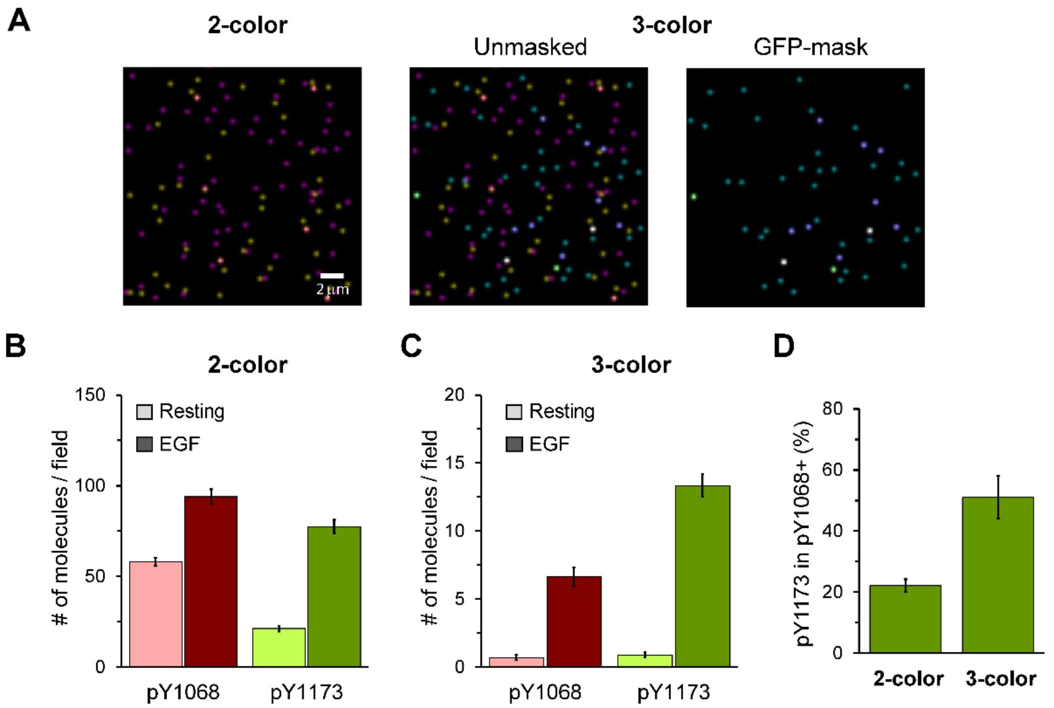


**Supplementary Figure S6. Importance of multi-color imaging for accurate quantification of phosphorylation percentages. (A)** Representative images displaying raw data and blob-reconstructed localized molecules from a 3-color SiMPull experiment. CHO-EGFR-GFP cells were stimulated with 25 nM EGF for 5 min at 37°C and assayed using anti-pY1068-CF555 (yellow) and anti-pY1173-CF640R (pink) antibodies. **(B)** Quantification of total number of pY1068 and pY1173 localizations per field of view when only those two channels are examined. EGFR-GFP channel was ignored for this quantification to emulate a 2-color SiMPull experiment. **(C)** Quantification of total number of pY1068 and pY1173 localizations per field of view using 3-color SiMPull. Here, the EGFR-GFP channel was used to identify pY1068 and pY1173 localizations overlapping with EGFR molecules, removing contributions from non-specific antibody binding. **(D)** In the absence of the EGFR-GFP channel to identify receptor locations, the 2-color SiMPull underestimates protein multi-phosphorylation. Number of receptors per condition, N>2400. Error bars are standard error of measured phosphorylation percentages.

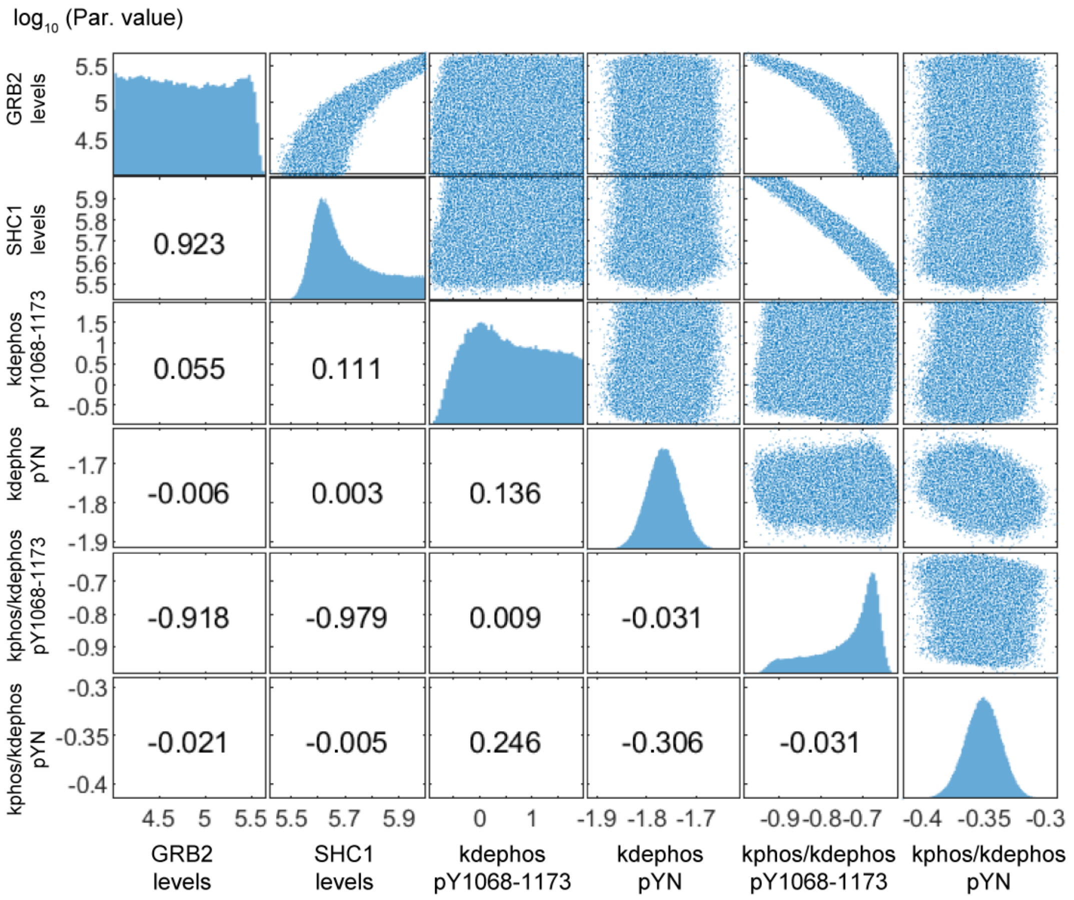


**Supplementary Figure S7: Quantification of parameter uncertainty by parallel tempering.** Results were obtained from >240,000 parameter sets sampled by parallel tempering (see Methods). The plots on the main diagonal show the sampled marginal probability distributions for each parameter. Scatter plots above the diagonal show the two-dimensional distributions of sampled parameter sets for each pair of parameters, to illustrate correlations between parameters. The plots below the diagonal contain the correlation coefficients (R values) between each parameter pair.

Reddy, Raven J., Aaron S. Gajadhar, Eric J. Swenson, Daniel A. Rothenberg, Timothy G. Curran, and Forest M. White. 2016. "Early Signaling Dynamics of the Epidermal Growth

Factor Receptor." *Proceedings of the National Academy of Sciences* 113 (11): 3114–19. https://doi.org/10.1073/pnas.1521288113.

Shi, Tujin, Mario Niepel, Jason E. McDermott, Yuqian Gao, Carrie D. Nicora, William B. Chrisler, Lye M. Markillie, et al. 2016. "Conservation of Protein Abundance Patterns Reveals the Regulatory Architecture of the EGFR-MAPK Pathway." *Science Signaling* 9 (436): rs6–rs6. https://doi.org/10.1126/scisignal.aaf0891.
