## Supplementary File S2 for "Integrating Multiplex SiMPull and Computational Modeling to Evaluate Combinatorial Aspects of EGFR Signaling": README.md

## Name
Salazar-Cavazos2019

## Reference
Salazar-Cavazos E, Nitta CF, Mitra ED, Wilson BS, Lidke KA, Hlavacek WS, Lidke DS. Integrating Multiplex SiMPull and Computational Modeling to Evaluate Combinatorial Aspects of EGFR Signaling. Under review.

## Description
A model of EGFR signaling that was parameterized using SiMPull measurements of multisite phosphorylation dynamics.

## Files
* [190127_CHO_EGFR_best-fit.bngl](190127_CHO_EGFR_best-fit.bngl): Model of EGFR signaling in CHO cells expressing EGFR-GFP, using the parameter values for best fit obtained through PyBNF.
* [190127_HeLa.bngl](190127_HeLa.bngl),  [190127_HMEC.bngl](190127_HMEC.bngl) and [190127_MCF10A.bngl](190127_MCF10A.bngl): Models of EGFR signaling for the differents cell lines. These models use as a base the model "190127\_CHO\_EGFR\_best-fit.bngl", with protein copy numbers for EGFR, Grb2 and Shc1 specific to each cell line.
* [190127_CHO_EGFR_sensitivity.bngl](190127_CHO_EGFR_sensitivity.bngl): Modification of the model "190127\_CHO\_EGFR\_best-fit.bngl" to perform sensitity analysis by increasing each of the parameter values in the model by 1%.
* [190127_CHO_EGFR_Epigen.bngl](190127_CHO_EGFR_Epigen.bngl): Model of EGFR signaling in CHO cells expressing EGFR-GFP, but in contrast to model "190127_CHO_EGFR_best-fit.bngl", it simulates EGFR activation with 20uM Epigen instead of 25nM EGF. The parameter value for EGFR dimer off-rate was varied to find the best fit for SiMPull results with Epigen ligand.
* [190127_CHO_HA_EGFR_L858R.bngl](190127_CHO_HA_EGFR_L858R.bngl): Model of EGFR signaling in CHO cells expressing HA-EGFR. The number of EGFR per cell was set to the experimentally estimated value for this cell line. The parameter value for kinase activity (phosphorylation rate) was varied to find best fit for experimental results with L858R mutant EGFR.
* [PyBNF-fitting-setup](PyBNF-fitting-setup): Files required to run parameterization of the base model in PyBioNetFit
  * [190127_CHO_EGFR_forBNF.bngl](PyBNF-fitting-setup/190127_CHO_EGFR_forBNF.bngl): Model file with unknown parameters
  * [dose_resp.exp](PyBNF-fitting-setup/dose_resp.exp), [EGF_25nM.exp](PyBNF-fitting-setup/EGF_25nM.exp): Experimental data files
  * [fit_v1_28.conf](PyBNF-fitting-setup/fit_v1_28.conf): PyBNF configuration file to perform fitting
  * [fit_bootstrap.conf](PyBNF-fitting-setup/fit_bootstrap.conf): PyBNF configuration file to perform bootstrapping
  * [fit_pt.conf](PyBNF-fitting-setup/fit_pt.conf): PyBNF configuration file to perform Bayesian uncertainty quantification by parallel tempering
  


## Notes
